# Genomic Signatures of Sexual Selection on Pollen-Expressed Genes in *Arabis alpina*

**DOI:** 10.1101/2021.09.02.457912

**Authors:** Juanita Gutiérrez-Valencia, Marco Fracassetti, Robert Horvath, Benjamin Laenen, Aurélie Désamore, Andreas D. Drouzas, Magne Friberg, Filip Kolář, Tanja Slotte

## Abstract

Fertilization in angiosperms involves the germination of pollen on the stigma, followed by the extrusion of a pollen tube that elongates through the style and delivers two sperm cells to the embryo sac. Sexual selection could occur throughout this process when male gametophytes compete for fertilization. The strength of sexual selection during pollen competition should be affected by the number of genotypes deposited on the stigma. As increased self-fertilization reduces the number of mating partners, and the genetic diversity and heterozygosity of populations, it should thereby reduce the intensity of sexual selection during pollen competition. Despite the prevalence of mating system shifts, few studies have directly compared the molecular signatures of sexual selection during pollen competition in populations with different mating systems. Here we analyzed whole-genome sequences from natural populations of *Arabis alpina*, a species showing mating system variation across its distribution, to test whether shifts from cross- to self-fertilization result in molecular signatures consistent with sexual selection on genes involved in pollen competition. We found evidence for efficient purifying selection on genes expressed in vegetative pollen, and overall weaker selection on sperm-expressed genes. This pattern was robust when controlling for gene expression level and specificity. In agreement with the expectation that sexual selection intensifies under cross-fertilization, we found that the efficacy of purifying selection on male gametophyte-expressed genes was significantly stronger in genetically more diverse and outbred populations. Our results show that intra-sexual competition shapes the evolution of pollen-expressed genes, and that its strength fades with increasing self-fertilization rates.

## Introduction

Sexual selection is a prevalent evolutionary force that impacts biological diversity in eukaryotes at all levels. This process favors the persistence of traits that increase an individual’s capacity to compete for fertilization opportunities, and manifests as two main mechanisms termed intrasexual competition (e.g. male-male contest) and intersexual selection (e.g. female choice) (Darwin 1871). Sexual selection is assumed to be significantly weaker in inbred lineages, such as those that have shifted from outcrossing to predominant self-fertilization (reviewed by Cutter 2019). This expectation is supported by the evolution of “selfing-syndromes” in plants, in which lineages that descend from outcrossing ancestors typically evolve a reduction or loss of traits otherwise involved in pollen transfer between individuals (Darwin 1876; Barrett 2002; Sicard and Lenhard 2011; Cutter 2019).

Sexual selection in flowering plants does not only affect the evolution of traits involved in pollen export. Stigmas normally receive pollen from different mating partners, and studies on seed paternity reflect the common occurrence of polyandry (Pannell and Labouche 2013). Such observations suggest that sexual selection might be an important evolutionary force shaping traits that mediate pollen-pistil interactions after pollination, including those affecting the performance of male gametophytes competing for fertilization (reviewed by Pannell and Labouche 2013; Tonnabel et al. 2021). Selfing could weaken the intensity of pollen competition through a reduction in the number of distinct mating partners (e.g. as under monogamy) (Lankinen et al. 2017), and by reducing the genetic diversity of the individuals in competition, which can affect overall reproductive success (Rowe and Houle 1996). However, our understanding of the consequences of the evolutionary transition to self-fertilization on pollen performance traits is still relatively limited (but see Hove and Mazer 2013; Lankinen et al. 2017; Mazer et al. 2018; Peters and Weis 2018; Harrison et al. 2019).

After pollen grains adhere to the stigma, they hydrate and germinate to grow a tubular extension that transports two genetically identical sperm cells through the style. Once this pollen tube reaches the female gametophyte, it bursts to deliver the sperm cells aided by the synergid cells. One of the sperm cells fuses with the egg giving rise to a diploid zygote, while simultaneously, the other sperm cell fertilizes the two nuclei of the central cell to form a triploid endosperm (Palanivelu and Tsukamoto 2012; Dresselhaus et al. 2016; Zheng et al. 2018). The sequence of events preceding fertilization creates the arena for pollen grains to compete based on their differential capacity to germinate (Austerlitz et al. 2012), grow their pollen tubes (Pasonen et al. 1999; Lankinen and Skogsmyr 2002), interfere with pollen from other donor plants (Varis et al. 2010) and navigate the transmitting tissue of the pistil in response to chemical cues (reviewed by Johnson et al. 2019). Experimental studies suggest that sexual selection is an important force driving the evolution of pollen traits (e.g. Skogsmyr and Lankinen 2002; Clark et al. 2006; Mazer et al. 2010; Lankinen et al. 2017). However, evolutionary genetic analyses suggest a minor role for sexual selection on genes expressed in sperm cells (e.g. Arunkumar et al. 2013). Interestingly, experiments show that sperm cells are disposable during the phase of pollen competition, as they are passively transported by pollen tubes and do not drive their own delivery (Zhang et al. 2017). Additionally, polyspermy avoidance mechanisms that ensure that only a single pair of sperm cells is delivered for double fertilization (reviewed by Dresselhaus and Franklin-Tong 2013) could result in reduced room for competition between sperm cells. Together, these findings reinforce the idea that sexual selection differentially impacts the individual components of male gametophytes.

What genetic signatures would intra-sexual competition produce on loci affecting pollen performance? If the fitness effects on pollen and the sporophyte are positively correlated, sexual selection would reduce the amount of genetic variation at such loci, as a result of both positive and purifying selection (Walsh and Charlesworth 1992). Theoretical work shows that standing genetic variation can persist if there is opposing selection pressures acting on the same allele in the haploid and diploid phase (Walsh and Charlesworth 1992; Immler et al. 2012), but such antagonistic pleiotropy should be more important if mutations that improve pollen performance are recessive and thus more frequently masked in the sporophyte (Peters and Weis, 2018). On the other hand, sexually selected traits can remain polymorphic despite directional selection if they are condition-dependent and if condition itself is highly genetically variable (Rowe and Houle 1996). Empirical results show that reproductive genes (e.g. those involved in gamete recognition, signaling and fertilization) evolve faster than genes expressed in non-reproductive tissues (as reviewed by Swanson and Vacquier, 2002; Clark et al. 2006). Although this pattern has been interpreted as a signal of adaptive evolution promoted by sexual selection, recent work has advocated for considering the contribution of relaxed purifying selection to elevated evolutionary rates (Dapper and Wade 2016; Dapper and Wade 2020). Indeed, recent genomic studies relying on polymorphism and divergence data have underscored the importance of quantifying both purifying and positive selection when studying evolutionary patterns and testing hypotheses on sexual selection at reproductive genes (e.g. Arunkumar et al. 2013; Gossmann et al. 2014; Harrison et al. 2019).

Factors other than sexual selection have also been proposed as important drivers of the evolution of reproductive genes in plants. First, in angiosperms, as many as 60% of genes are expressed in the haploid gametophytic phase, and about 7-11% of genes are exclusively expressed at that stage (Honys and Twell 2004; Pina et al. 2005; Borges et al. 2008). In contrast to diploid tissues, recessive and partially recessive new mutations arising in genes expressed in haploid stages are directly exposed to the action of selection, which should aid the removal of deleterious mutations and fix the advantageous ones (Haldane 1932; Haldane 1933; Kondrashov and Crow 1991; Charlesworth and Charlesworth 1992; Gerstein and Otto 2009). Second, expression-related variables are known to impact rates of protein evolution. In general, broadly and highly expressed genes show slower amino acid substitution rates compared to tissue-specific or lowly expressed genes (Wright et al. 2004; Drummond et al. 2005; Slotte et al. 2011; Yang and Gaut 2011; Zhang and Yang 2015). The transcriptomes of haploid cells tend to be enriched in specifically-expressed genes, which is thought to reduce the evolutionary constraints acting on such genes (Szövényi et al. 2013). For instance, low expression levels in sperm cells might explain the accumulation of slightly deleterious mutations in sperm-expressed genes (Arunkumar et al. 2013). Hence, it is necessary to investigate the potential contribution of transcript specificity and abundance to molecular evolution when studying signatures of selection on gametophyte-expressed genes.

To more directly understand the contribution of sexual selection to the evolution of male gametophyte-expressed genes, it is necessary to investigate if variation in the strength of sexual selection alters the fate of new mutations in these genes. If sexual selection is important to the evolution of pollen expressed genes, alleles promoting faster germination and tube growth rates should be selected for, and alleles negatively affecting these traits should be removed (Mazer et al. 2010). Elevated selfing rates are expected to lead to reduced intensity of sexual selection at the pollen stage (Mazer et al. 2010). This outcome could be further magnified in selfing populations that experience a reduced efficacy of selection genome-wide due to reductions in the effective population size and the increased impact of linked selection (Wright et al. 2013; Slotte 2014; Hartfield et al. 2017).

Experimental studies have found conflicting results regarding the impact of mating system shifts on the rates at which pollen grains germinate and grow their pollen tubes (e.g. Smith-Huerta 1996; Taylor and Williams 2012; Hove and Mazer 2013; Lankinen et al. 2017), and the impact it might have on the offspring (Willi 2013). These inconsistencies could in part be explained by methodological differences between experiments (as discussed by Mazer et al. 2018). Recently, population genomic studies have started to bring further insight into the effects of mating system variation by characterizing the signatures of selection on genes involved in pollen competition. For instance, using divergence and polymorphism-based analyses, Harrison et al. (2019) found that pollen-expressed genes in *Arabidopsis thaliana* experienced a relaxation in selection in association with the transition to predominantly self-fertilization. However, it remains unknown how general these findings are and how rapidly such effects may occur. Direct within-species studies comparing genomic signatures of selection between outcrossing and selfing populations could bring further light on this matter, and could be more robust to confounding effects present in between-species comparisons.

In this study, we aimed to compare signatures of selection on genes expressed in male and female gametophytes, across populations with different levels of genetic diversity and inbreeding rates. Specifically, we wanted to test if differences in selection can be attributed to the intensity of sexual selection that populations with different mating systems experience. For this purpose, we analyzed whole-genome sequences from natural populations of *Arabis alpina* with different mating systems. This arctic-alpine perennial herb is ideally suited for this purpose, because different populations show contrasting inbreeding levels across the species’ distribution, as a joint consequence of demographic history and mating system changes (Ansell et al. 2008; Tedder et al. 2011; Laenen et al. 2018). Previous experimental studies have shown that populations of *A. alpina* have evolved a wide spectrum of mating system strategies ranging from self-incompatibility and obligate outcrossing, to self-compatibility dominating most or the entire population (Tedder et al. 2011; Toräng et al. 2017; Petrén et al. 2021). It currently remains unknown whether loss of self-incompatibility has occurred several times independently in this species, but all self-compatible populations studied in detail exhibit a selfing syndrome both in terms of a reduced floral size (Tedder et al. 2011; Petrén et al. 2021) and a reduced floral scent signal (Petrén et al. 2021).

Here, we first contrast the prevalence of purifying and positive selection in an outcrossing *A. alpina* population, focusing on genes expressed in male and female gametophytes and sporophytes. Then, we investigate the impact of potential confounding variables such as cell/tissue ploidy and expression specificity and abundance on signatures of selection. Finally, we test if purifying selection on genes expressed in gametophyte components varies with inbreeding and polymorphism levels across *A. alpina* populations with different mating systems. We rely primarily on the uniquely detailed and replicated expression data from *A. thaliana* for identification of genes expressed in male and female gametophytic components and we show that our main results are robust to this choice. Taken together, our results show that selection differentially impacts vegetative pollen cell and sperm-expressed genes, an observation that is robust to controlling for differences in gene expression breadth and abundance. We further found that purifying selection on genes expressed in vegetative pollen, pollen tubes and sperm was specifically more efficient in more polymorphic, outbred and likely outcrossing populations, as expected under efficient sexual selection.

## Results

### Efficient Purifying Selection on Gametophyte-expressed Genes

To test for a molecular signature of sexual selection on genes expressed in gametophytes, we generated whole-genome sequences of 20 individuals from an outcrossing (mean inbreeding coefficient *F_IS_*=1.8×10^−2^) and genetically polymorphic (*π_S_*=0.006) population of *A. alpina* from Greece (population Gre2) where sexual selection would be expected to be efficient (supplementary figs 1-3, supplementary table 1, Supplementary Material). Using polymorphism data from this population and divergence from two closely related outgroup species at nonsynonymous (only 0-fold degenerate sites considered) and synonymous sites (only 4-fold degenerate sites considered), we estimated the impact of selection on sets of all genes identified as expressed in the male gametophyte (12,398 genes). For comparison, we conducted the same analyses with all genes expressed in the female gametophyte (10,893 genes) and in the sporophyte (10,000 randomly selected genes identified as expressed in the vegetative shoot apex, leaf, petiole and/or root). These gene sets were identified using expression data from *A. thaliana*, followed by the determination of the corresponding *A. alpina* orthologs (see Materials and Methods for details).

Analyses of the distribution of negative fitness effects (DFE) of new nonsynonymous mutations showed that genes expressed in gametophytes had a lower fraction of nearly neutral new nonsynonymous mutations (0 < *N_e_s* < 1) and are thus under stronger purifying selection than sporophyte-expressed genes in our outcrossing *A. alpina* population (Kruskal-Wallis test followed by Dunn’s test with Bonferroni corrected *P*-values, *P*<0.001, based on contrasts of 200 bootstrap replicates; see Materials and Methods for details) (fig. 1). Genes expressed in female gametophytes were under stronger purifying selection than those expressed in male gametophytes (*P*<0.001, Kruskal-Wallis test followed by Dunn’s test with Bonferroni correction, based on contrasts of 200 bootstrap replicates), with 13.17±0.14% and 14.91±0.09% of nearly neutrally-evolving mutations (0 < *N_e_s* < 1), respectively. Estimates of the proportion of adaptive nonsynonymous substitutions (*α*) suggest more frequent adaptation in male and female gametophyte-expressed genes compared to sporophyte-expressed genes (male=0.074±0.012, female=0.082±0.012, sporophyte=0.064±0.007, *P*<0.001, Kruskal-Wallis test followed by Dunn’s test with Bonferroni correction, based on contrasts of 200 bootstrap replicates). However, there were no significant differences between gene sets in the proportion of adaptive nonsynonymous substitutions relative to synonymous substitutions (*ω_α_*) (male=0.011±0.006, female=0.011±0.010, sporophyte=0.010± 0.001, *P*>0.05, Kruskal-Wallis test followed by Dunn’s test with Bonferroni correction, based on contrasts of 200 bootstrap replicates) (supplementary fig. 4, Supplementary Material). In agreement with the DFE results, gametophyte-expressed genes showed a lower ratio of nonsynonymous to synonymous polymorphism (*π_N_/π_S_*) than sporophyte-expressed genes *(π_N_/π_S_* male gametophyte=0.17, female gametophyte=0.16 and sporophyte=0.18), and we estimated a higher ratio of non-synonymous to synonymous substitutions in sporophyte-expressed than in gametophyte-expressed genes (*d_N_/d_S_* male gametophyte=0.15, female gametophyte=0.13 and sporophyte=0.16) (supplementary table 2, Supplementary Material).

**Figure 1.**
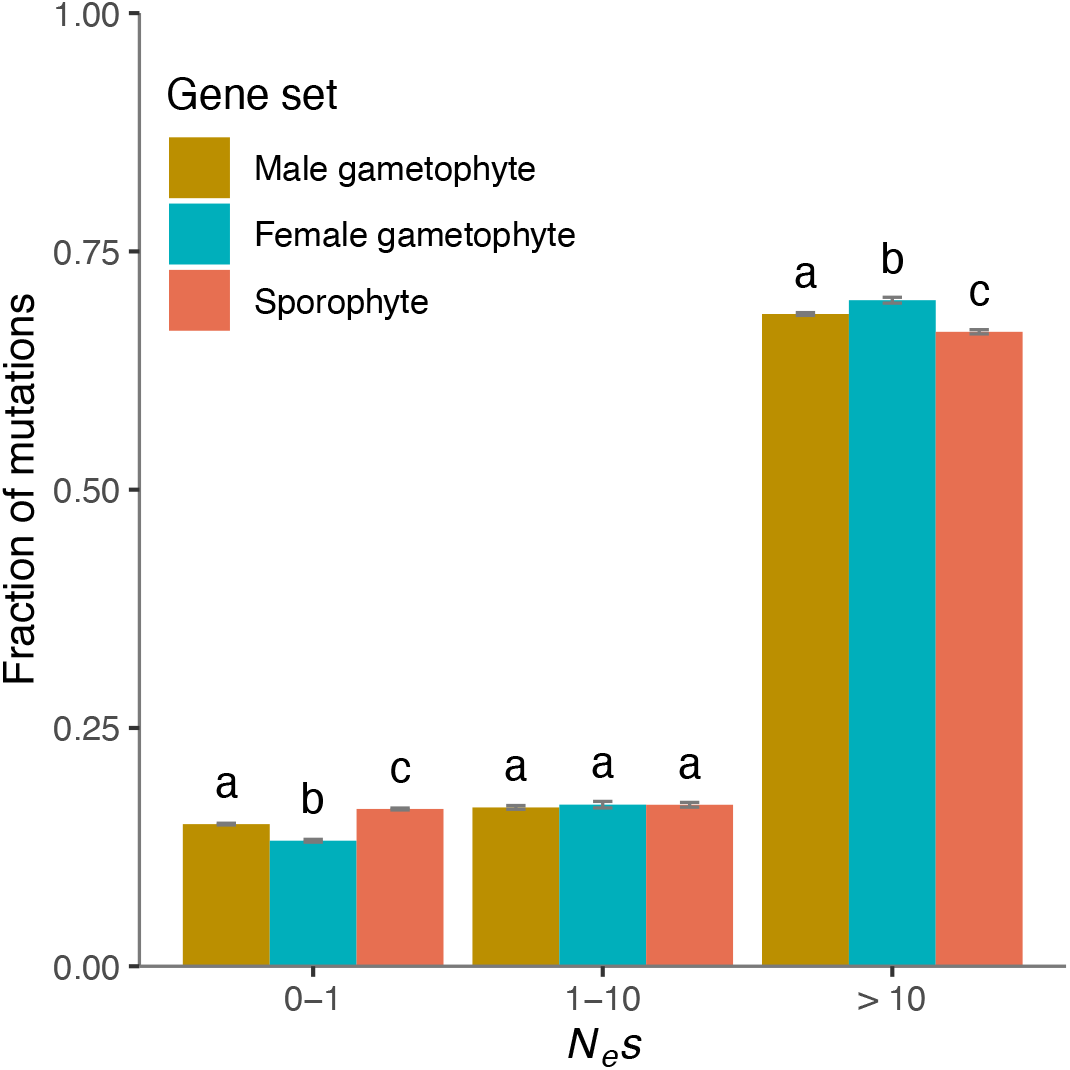
Comparison of DFE estimates for genes expressed in the male gametophyte (*n*=12,398), the female gametophyte (*n*=10,893) and sporophytic tissues (*n*=10,000, random sample) for the outcrossing population Gre2. Error bars represent the bootstrap standard error (200 replicates). Different letters denote statistically significant differences between gene set categories.

### Haploid Selection Cannot Explain the Impact of Purifying Selection on Gametophytic Genes

Stronger purifying selection on gametophyte-expressed genes could either be a result of stronger functional constraints on genes involved in reproductive functions, or a result of haploid selection. To assess whether differences in ploidy level shape selection on genes expressed in gametophytes, we estimated DFE, *α* and *ω_α_* for all genes expressed in each of the components of the male gametophyte (vegetative pollen cell, pollen tube and sperm cells) (fig. 2) and the female gametophyte (synergids, egg and nucellus), and compared them to the diploid nucellar tissue and unpollinated pistils. By analyzing the components of the female gametophyte individually, we aimed not only to distinguish the patterns of selection affecting cell types with different ploidy, but also to include a broader representation of structures involved in reproduction as a control. If genes expressed in haploid cell types/tissues are consistently more impacted by purifying and positive selection than genes expressed in the two diploid tissues, that would suggest that higher efficiency of selection in gametophytes is primarily explained by haploid selection. We considered genes expressed in the vegetative cell of pollen and pollen tubes as different categories because they represent different stages of pollen competition (Lankinen and Karlsson Green 2015), although the pollen tube is derived from the vegetative cell of pollen. The gene sets expressed in the vegetative pollen cell that we analyze here are involved in functions such as signaling, vesicle trafficking, cell wall metabolism, transport across membranes, and the cytoskeleton modification, all of which are associated with the process of pollen germination on the surface of the stigma (Pina et al. 2005, Borges et al. 2008). In contrast, the gene set expressed in semi-*in vivo* grown pollen tubes has been linked to the functions of protein recognition, signaling, transmembrane receptor activity and defense response (Qin et al. 2009), likely involved in the perception and response to pistil guidance and recognition between mating partners, as well as male-male competition.

**Figure 2.**
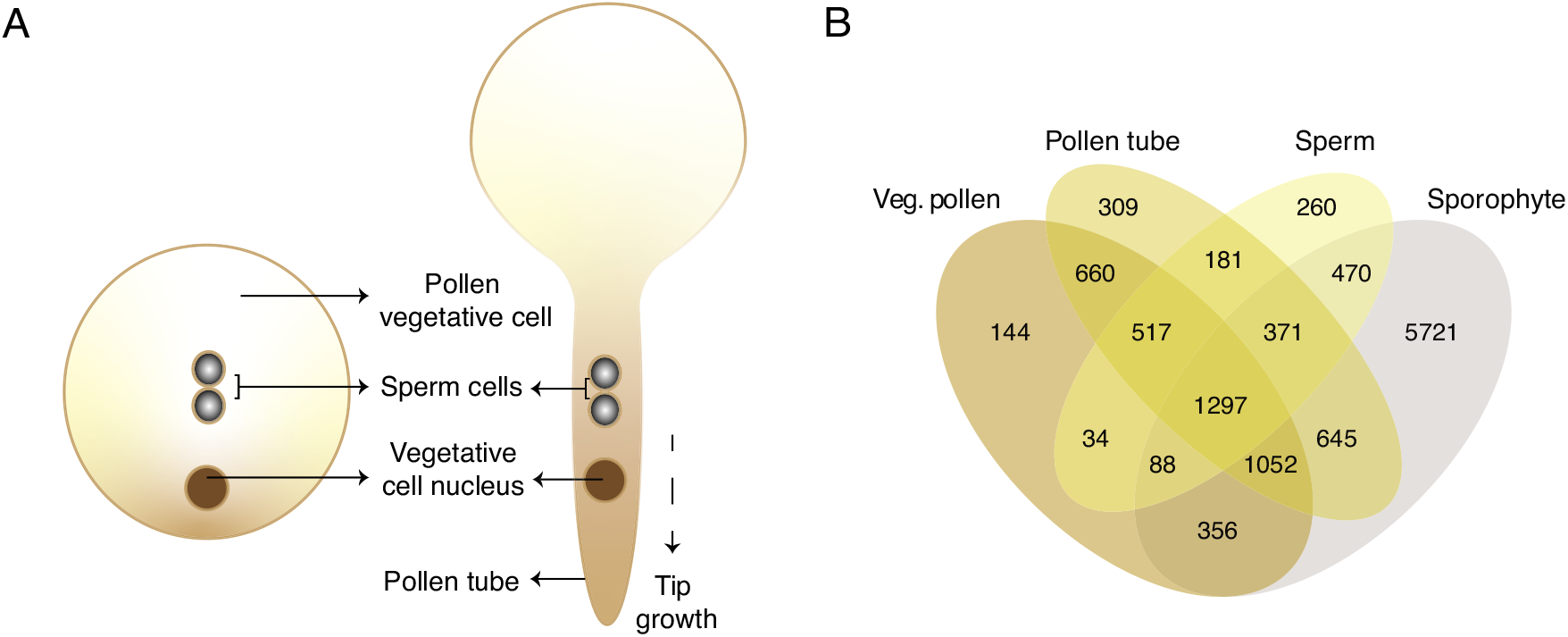
(**A**) Diagram of the haploid male gametophyte and its individual components, comprising the vegetative components of mature vegetative pollen (pollen vegetative cell and its nucleus) and two non-motile sperm cells. After landing on the stigma, the membrane of the vegetative cell protrudes forming the pollen tubes that prolongs through the pistil transmitting tissue, to finally burst and relase the two sperm cells to the female gametophyte wherein double fertilization occurs. (**B**) Venn diagram showing the number of genes expressed in the vegetative components of vegetative pollen, pollen tubes and sperm (for details, see Materials and Methods).

We found that purifying selection was not systematically stronger for genes expressed in haploid components compared to those expressed in the diploid nucellus and unpollinated pistils. Interestingly, the DFE differed significantly between the different haploid gene sets (fig. 3a). In agreement with the first analysis, we found that genes expressed in components of the male gametophyte are less constrained than those expressed in female components. There were significant differences in the proportion of nearly neutral new nonsynonymous mutations (0 < *N_e_s* <1) between different haploid components of the male gametophyte (figs. 2a and 3a), with genes expressed in pollen tubes being under weaker purifying selection than those expressed in vegetative pollen cells (0 < *N_e_s* < 1: 14.97±0.14% vs 13.67±0.14%, respectively, *P*<0.001, Kruskal-Wallis test followed by Dunn’s test with Bonferroni correction, based on contrasts of 200 bootstrap replicates). Genes expressed in sperm cells showed the greatest proportion of new nearly neutral nonsynonymous mutations (16.33±0.19%, *P*<0.001 for all pairwise tests, Kruskal-Wallis test followed by Dunn’s test with Bonferroni correction, based on contrasts of 200 bootstrap replicates), and *π_N_/π_S_* and *d_N_/d_S_* estimates were also highest for sperm-expressed genes (*π_N_/π_S_* vegetative pollen=0.16, pollen tubes=0.18 and sperm=0.19; *d_N_d_S_* vegetative pollen=0.14, pollen tube=0.15 and sperm=0.16) (supplementary table 2, Supplementary Material).

**Figure 3.**
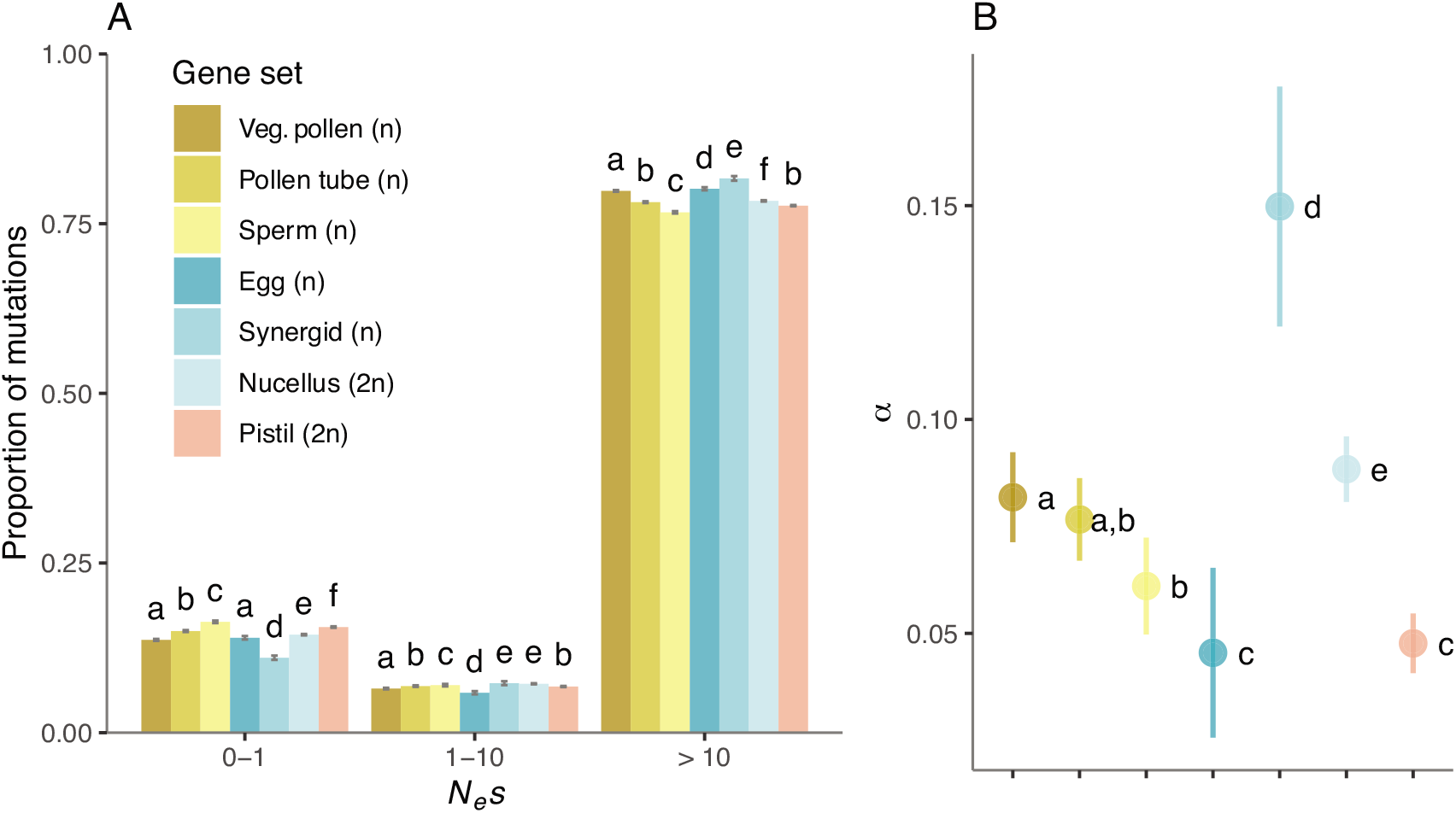
Comparison of the estimates of (**A**) the distribution of negative fitness effects (DFE) and (**B**) the proportion of adaptive substitutions (*α*) for genes expressed in individual haploid components of the male and female gametophytes and the diploid nucellus. Number of genes in each set: vegetative pollen (*n*=4,148), sperm (*n*=3,218), pollen tubes (*n*=5,032), synergids (*n*=511), egg (*n*=1,607), nucellus (*n*=6,875) and unpollinated pistil (*n*= 9,555). Error bars represent the bootstrap standard error (200 replicates). Different letters denote statistically significant differences between gene sets.

We did not find evidence for consistently higher rates of adaptation in genes expressed in haploid cells compared to genes expressed in diploid tissues (fig. 3b, supplementary table 2, supplementary fig. 5a, Supplementary Material). Among male gametophyte components, genes expressed in vegetative pollen cells were more impacted by positive selection than those expressed in sperm cells (*α*=0.082± 0.011 and 0.061± 0.011 respectively, *P*<0.001; *ω_α_*=0.011± 0.001 and 0.009±0.002 respectively, *P*<0.001, Kruskal-Wallis test followed by Dunn’s test with Bonferroni correction, based on contrasts of 200 bootstrap replicates). Positive selection on genes expressed in pollen tubes did not differ significantly from those two gene sets (*α*=0.077±0.010 and *ω_α_*=0.011±0.001, NS, Dunn’s test based on contrasts of 200 bootstrap replicates, Bonferroni corrected *P*-values). Genes expressed in haploid synergids were the most affected by positive selection (*α*=0.150±0.028 and *ω_α_*=0.017±0.003, *P*<0.001, Kruskal-Wallis test followed by Dunn’s test with Bonferroni correction, based on contrasts of 200 bootstrap replicates), followed by the diploid nucellus tissue (*α*=0.088±0.008 and *ω_α_*=0.013±0.001), and finally diploid pistils and haploid egg cells (*α*=0.048±0.010 and *ω_α_*=0.007±0.001, and *α*=0.045±0.020 and *ω_α_*=0.006±0.003 respectively). Our results suggest that signatures of selection observed in genes expressed in gametophytes cannot be attributed completely to their haploid condition. Moreover, selection differentially affects genes expressed in haploid condition in vegetative pollen, pollen tubes and sperm.

### Gene Expression Variables Do Not Drive Contrasting Patterns of Selection on Genes Expressed in Vegetative Pollen and Sperm Cells

If differences in selection on genes expressed in vegetative pollen, pollen tubes and sperm are caused by intrinsic expression disparities between sets of genes, these differences should not persist after controlling for these variables. To control for expression specificity, we reduced our gene sets to only include those genes that were exclusively expressed in each component of the male gametophyte. We intersected the original sets of genes expressed in vegetative pollen, pollen tubes and sperm with a set of genes expressed in the sporophyte (10,000 randomly selected genes expressed in the shoot apex, leaf, petiole and/or root) (fig. 2b), and then repeated our analyses of purifying and positive selection on reduced gene sets. We identified 144 genes putatively exclusive to vegetative pollen cells (3.47% of the initial set), 309 in pollen tubes (6.14%), 260 in sperm (8.08%) and 5,721 in sporophyte tissues (57.21%) (fig. 2b). In agreement with estimates based on the full gene sets, genes expressed exclusively in vegetative pollen cells experienced stronger purifying selection than those exclusively expressed in pollen tubes, while sperm cells were under significantly weaker selection (fig. 4a). Based on exclusively expressed gene sets, we again found no evidence for consistently stronger purifying selection on genes expressed in haploid components (supplementary fig. 7a, Supplementary Material). To make our results more comparable to a previous study conducted on *Capsella grandiflora* (Arunkumar et al. 2013), we repeated this analysis with exclusively expressed genes in the components of male gametophytes identified after intersecting the initial sets with data on seedling-expressed genes. As before, we inferred that vegetative pollen-expressed genes (2.24% of the initial set) experienced the strongest intensity of purifying selection, and sperm-expressed genes (6.80% of the initial set) experienced significantly weaker purifying selection (supplementary fig. 7b, Supplementary Material).

**Figure 4.**
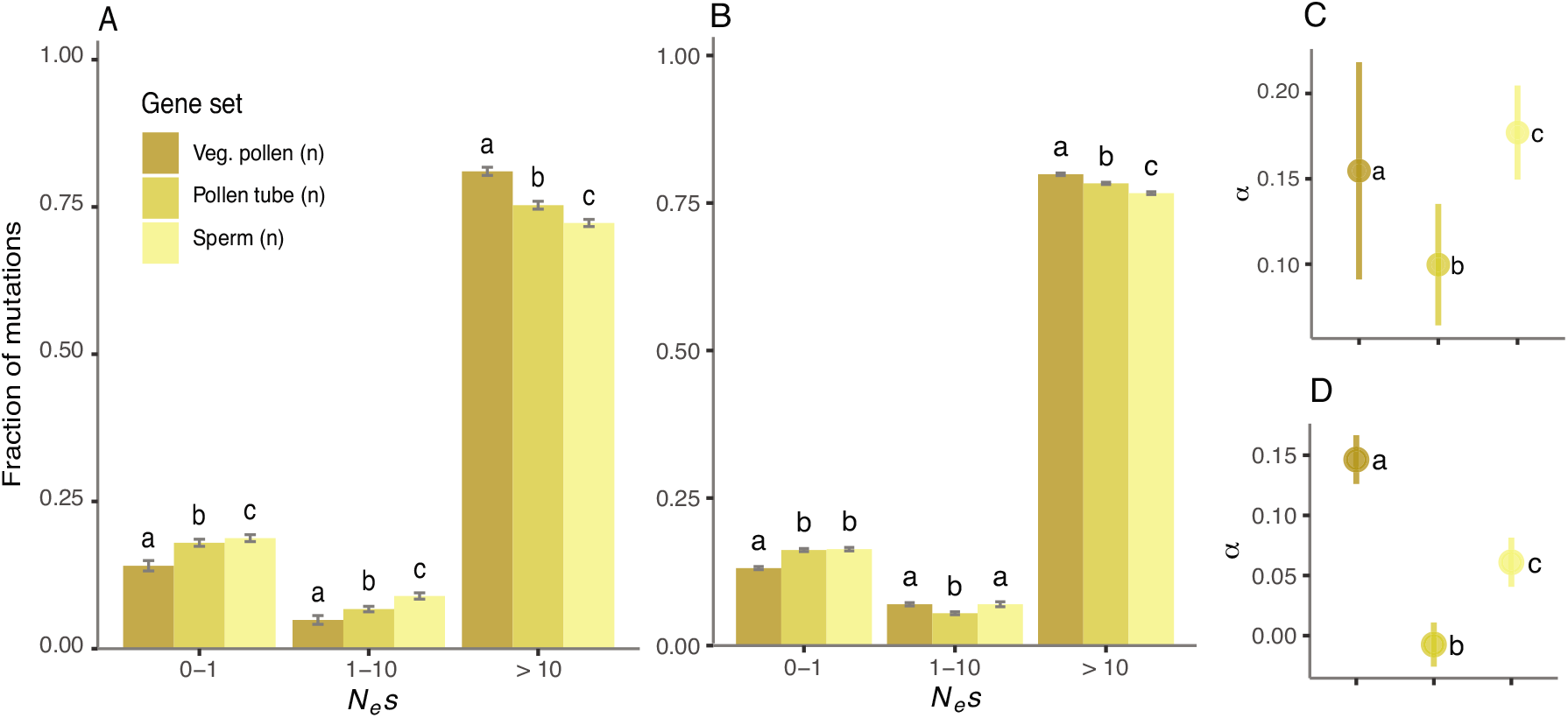
Comparison of the distribution of negative fitness effects (DFE) (**A, B**) and the proportion of adaptive substitutions (*α*) (**C, D**) after controlling for expression variables. **A, C.** Control for expression specificity: vegetative pollen (*n*=144), pollen tubes (*n*=309) and sperm cells (*n*=260). **B, D.** Control for transcript abundance: vegetative pollen (*n*=1,552), pollen tubes (*n*=2,387) and sperm (*n*=3,218). Error bars represent the bootstrap standard error (200 replicates). Different letters denote statistically significant differences between gene sets.

We controlled for expression level using a matched comparison group approach. Briefly, we reduced each original data set to a subset of genes matching the distribution of transcript abundances for the component of male gametophytes with the fewest expressed genes, i.e. sperm cells (n=3218 genes, median=834) (see Materials and Methods for details). This resulted in a total of 1,552 genes expressed in vegetative pollen (37% of the initial set of genes, median before correction=1092, median after correction=837) and 2,387 expressed in pollen tubes (47% of the initial set of genes, median before correction=862, median after correction=844) (supplementary fig. 8, Supplementary material). Again, we found that vegetative pollen cell-expressed genes experienced the strongest levels of purifying selection, followed by genes expressed in pollen tubes and sperm cells (fig. 4b). Thus, differences in the DFE between genes expressed in vegetative pollen, pollen tubes and sperm cells were robust to controls for expression level and specificity.

Positive selection was significantly more prevalent in genes expressed in vegetative pollen compared with other components of the male gametophyte after controlling for expression abundance, but not after accounting for gene expression breadth. When controlling for differences in transcript abundance, vegetative pollen-expressed genes still showed the highest rates of adaptive evolution (*α*=0.14±0.02), followed by genes expressed in sperm (*α*=0.06±0.02) and pollen tubes (*α*=−0.01±0.02) (fig. 4d). These results held for *ω_α_* estimates as well (supplementary fig. 5c, Supplementary Material). However, when restricting our analyses to sets of exclusively expressed genes, we estimated the proportion of adaptive substitutions to be significantly higher in sperm cells (*α*=0.18±0.03) compared to vegetative pollen cells (*α*=0.15±0.06) and pollen tubes (*α*=0.10±0.03) and (fig. 4c). The same pattern was observed for estimates of *ω_α_* (supplementary fig. 5b, Supplementary Material).

We further tested how sensitive our results were to our reliance on expression data from *A. thaliana* to identify pollen-expressed genes. As no similarly detailed or replicated expression resource was available for *A. alpina* or other Brassicaceae species, we identified pollen- and unpollinated pistil-expressed genes based on publicly available data from *Brassica napus* (Lohani et al. 2021) and *A. thaliana* (see Materials and Methods and Supplementary Note 1 for details). We inferred the DFE of the *A. alpina* orthologs of these gene sets and inferred stronger purifying selection on pollen than on pistil-expressed genes in the Gre2 population, both based on *B. napus* and *A. thaliana* data (0 < *N_e_s* < 1 estimates based on *A. thaliana* expression data= pollen: 13.67±0.14%, pistil: 15.56±0.10%; B. *napus*= pollen: 12.03±0.17%, pistil: 13.97±0.09%). We also found no significant differences in adaptive evolution of pollen-expressed genes, but for pistil-expressed genes *A. thaliana-based* estimates were lower (supplementary fig. 6a, Supplementary Material). Estimates of purifying but not necessarily positive selection thus appear to be robust to our strategy to identify pollen and pistil-expressed genes (supplementary fig. 6a, Supplementary Note 1, Supplementary Material).

Our results suggest that despite differences in the ubiquity and abundance of genes expressed, vegetative pollen cell and sperm-expressed genes show contrasting patterns of purifying selection, with vegetative pollen-expressed genes consistently showing a lower proportion of effectively neutral sites. In contrast, gene expression breadth seems to affect the differences in adaptive evolution between vegetative pollen and sperm. These results show that in a polymorphic and outbred population, contrasting signatures of purifying selection in vegetative pollen- and sperm-expressed genes are not caused by the higher tissue specificity and lower expression levels of genes expressed in sperm, allowing us to further investigate how these sets of genes are impacted by different intensities of intrasexual competition.

### Weaker Purifying Selection on Pollen-Expressed Genes in More Inbred and Less Polymorphic Populations

We hypothesized that in populations with low levels of polymorphism and high inbreeding rates, pollen-expressed genes would accumulate a higher proportion of slightly deleterious mutations due to relaxed sexual selection. To test this hypothesis, we generated and analyzed whole-genome sequences from 228 individuals from 13 populations of *A. alpina*, with different mating systems and demographic histories (Laenen et al. 2018) (supplementary fig. 9a, supplementary table 3, Supplementary Material). We refrained from comparing *α* and *ω_α_* across populations because those estimates would rely on the same divergence data (see Materials and Methods) and would therefore not be independent.

To characterize levels of genetic diversity and inbreeding we first calculated and compared nucleotide diversity (*π*), and two estimates of the inbreeding coefficient based on deviations between observed and expected genotype frequencies (*F_IS_*) and based on runs of homozygosity genome-wide (*F*_ROH_) for each population (supplementary figs. 2 and 3, supplementary table 1, Supplementary Material). We found that the most polymorphic and outbred populations came from Greece and Italy, while populations from central and eastern Europe had lower polymorphism levels and variable levels of inbreeding, and the lowest levels of polymorphism were found in highly selfing and inbred Scandinavian populations (supplementary table 1, Supplementary Material). Overall, population structure was largely concordant with geographic sampling locality, suggesting that *F_IS_* should not be greatly overestimated due to the Wahlund effect (supplementary fig. 9b, Supplementary Material). Altogether, estimates of *π* at 4-fold synonymous sites spanned from 1.1×10^−5^ to 6.2×10^−3^, whereas mean *F_IS_* values for 4-fold synonymous sites ranged from 0.02±0.05 to 0.97±0.08, and mean *F*_ROH_ (ROH > 500 Kb) ranged from 0.23±0.04 to 0.92±0.01 (supplementary figs. 2, 3 and 10, Supplementary Material). Previous work has shown that *F_IS_* estimates are consistent with the functioning of the self-incompatibility system in *A. alpina* (Tedder et al. 2011), suggesting it could provide a proxy for classification of mating system. Our estimates of *F_IS_* and *F*_ROH_ (supplementary figs. 2 and 3, Supplementary Material) allowed us to tentatively classify populations in two discrete categories: predominantly outcrossing populations with a mean 4-fold *F_IS_* < 0.04 corresponding to a selfing rate of < 0.08 assuming equilibrium, and predominantly selfing populations with a mean 4-fold *F_IS_* > 0.85, corresponding to a selfing rate under equilibrium of > 0.92 (See Materials and Methods for details). This data set therefore offers a wide range of variation in terms of polymorphism level and degree of inbreeding to test for the impact of sexual selection.

One potential complication when testing for an effect of diversity and inbreeding on sexual selection is that strong reductions in the effective population size, e.g. due to demographic bottlenecks and/or linked selection, can result in reduced efficacy of selection genome-wide (Wright et al. 2013; Mattila et al. 2020). In such populations, it can be difficult to specifically attribute relaxed purifying selection on gametophyte-expressed genes to relaxed sexual selection. To assess whether some of our populations exhibited a signature of genome-wide relaxed selection, we first estimated DFE in a set of 10,000 randomly selected genome-wide genes in all populations. We found that the three populations with the lowest *π* at 4-fold sites, including two Scandinavian populations, appear to have experienced such an effect, with most (or even all) analyzed sites evolving either as effectively neutral or under weak purifying selection (supplementary fig. 11, Supplementary Material). We therefore excluded these populations from further consideration in our analyses of gametophyte-expressed genes.

Next, we asked whether the efficacy of purifying selection on genes expressed in the male gametophyte varies with nucleotide diversity and inbreeding level in the remaining ten populations. We found that *π* at 4-fold synonymous sites was significantly negatively correlated with the fraction of nearly neutral new nonsynonymous mutations for genes expressed in vegetative pollen, pollen tubes and sperm, as expected if selection on these genes were stronger in more polymorphic populations (fig. 5a-c, table 1, supplementary fig. 10e-g, Supplementary Material). Although significant, this negative correlation was weaker for the analyses based on genome-wide estimates (10,000 randomly selected genes) (fig. 5d, table 1, supplementary fig. 10h, Supplementary Material). After correcting for phylogenetic non-independence among populations, these results held true for vegetative pollen, pollen tube and sperm-expressed genes, but not for genome-wide estimates (table 1, supplementary fig. 12, Supplementary Material). These findings show that purifying selection on genes expressed in vegetative pollen, pollen tubes and sperm is weaker in less polymorphic populations, and that this result cannot be fully explained by a genome-wide relaxation of purifying selection in less polymorphic populations.

**Figure 5.**
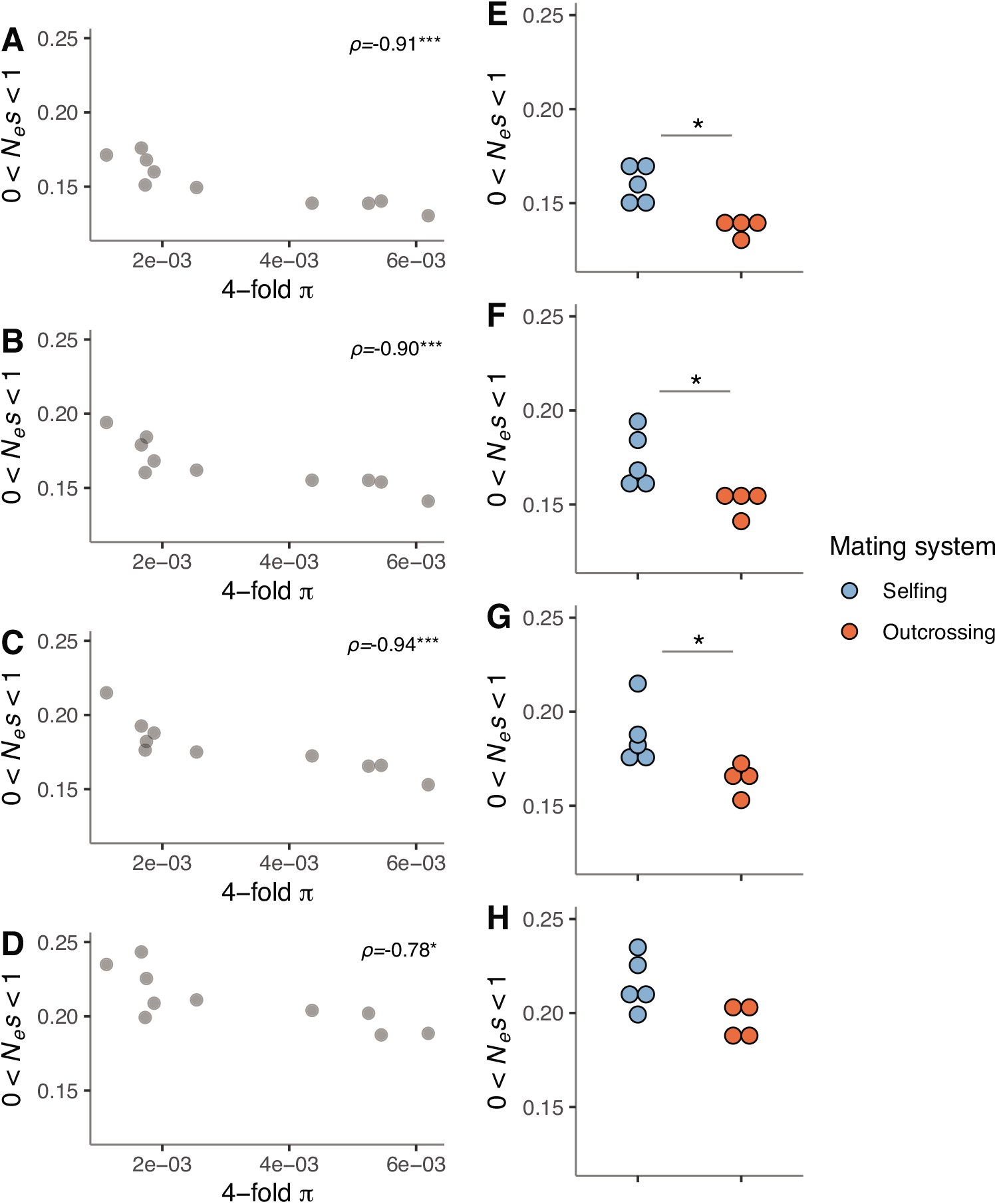
(**A**-**D**) Scatterplots depicting the correlation between estimates of 4-fold *π* and the fraction of nearly neutral sites (0 < *N_e_s* < 1) across *A. alpina* populations (*n*=10). Spearman’s rank correlation rho (*ρ*) values and the statistical significance of the association between these variables are shown for (**A**) vegetative pollen, (**B**) pollen tubes, (**C**) sperm and (**D**) 10,000 genome-wide randomly selected genes (*: *P*<0.05, **: *P*<0.01, ***: *P*<0.001). (**E**-**H**) Comparison of the estimates of the fraction of nearly neutral sites between predominantly outcrossing (*n*=4) and selfing (*n*=5) populations for each data set: (**E**) vegetative pollen, (**F**) pollen tubes, (**G**) sperm and (**H**) 10,000 genome-wide randomly selected genes (*: *P*<0.05, Wilcoxon rank-sum test).

**Table 1.** Estimates of Spearman’s rank correlation coefficient (*ρ*) between 4-fold *π* values and the fraction of nearly neutral new 0-fold degenerate mutations (DFE category of 0 < *N_e_s* < 1) before and after phylogenetically independent contrast (PIC) correction for each set of genes expressed in the components of male gametophytes and a set of 10,000 randomly selected genes representing genome-wide patterns.

| Cell type / tissue | Before PIC correction |  | After PIC correction |  |
| --- | --- | --- | --- | --- |
| | $\rho$ | *P*-value | $\rho$ | *P*-value |
| **Vegetative pollen cell** | -0.91 | 0.0005 | -0.98 | 0.0001 |
| **Pollen tube** | -0.90 | 0.0009 | -0.87 | 0.0045 |
| **Sperm** | -0.94 | 0.0001 | -0.83 | 0.0083 |
| **Genome-wide** | -0.78 | 0.0116 | -0.67 | 0.0589 |

We further investigated if, as in the case of polymorphism, inbreeding impacts purifying selection on male gametophyte-expressed genes. For each analyzed set of genes, we tested if the proportion of nearly neutral new non-synonymous mutations differed between predominantly outcrossing and selfing populations. We found that predominantly outcrossing populations show a significantly lower fraction of nearly neutral new non-synonymous mutations compared to the predominantly selfing ones for genes expressed in vegetative pollen, pollen tubes and sperm (*P*<0.05 in all cases, Wilcoxon rank-sum test), but not for sporophyte-expressed genes (*P*=0.07, Wilcoxon rank-sum test) (fig. 5e-h). Taken together, these results suggest that purifying selection at male gametophyte-expressed genes is weaker in the predominantly selfing populations than in the outcrossing populations.

### Contrasting Patterns of Selection on Vegetative Pollen and Sperm-Expressed Genes Across Populations

Our previous analyses showed a consistent pattern of weaker selection on sperm-expressed than vegetative pollen-expressed genes. We asked whether this pattern was exclusive to outbred populations, or whether genes expressed in vegetative pollen cells were generally under stronger constraint than those expressed in sperm cells regardless of population. Thus, we estimated DFE for genes expressed in these two components of the male gametophyte and genome-wide estimates, and compared the inferred fraction of nearly neutral new non-synonymous mutations (0 < *N_e_s* < 1) across populations. Regardless of population level of polymorphism or inbreeding, purifying selection was always stronger on genes expressed in vegetative pollen cells than in sperm cells, and than genome-wide estimates. These results suggest that there are consistently stronger relative constraints acting on vegetative pollen-expressed genes than on sperm-expressed genes (supplementary fig. 13, Supplementary Material).

### Genetic Polymorphism does not Correlate with the Proportion of Effectively Neutral Sites in Synergid-Expressed Genes

Among all the individual components of the male and female gametophytes, synergid-expressed genes show the strongest levels of purifying selection (fig. 3a). We investigated whether, as in the case of genes expressed in the male gametophyte, the strength of purifying selection would correlate with levels of genetic polymorphism. Contrary to this expectation, we found no correlation between 4-fold synonymous *π* and the fraction of nearly neutral new nonsynonymous mutations for this gene set (supplementary fig. 14, Supplementary Material).

## Discussion

### Prevalent Signatures of Purifying Selection on Vegetative Pollen-Expressed Genes

It has long been proposed that sexual selection is an important driving force for the evolution of genes expressed in pollen (Bernasconi et al. 2004; Mazer et al. 2010; Lankinen and Karlsson Green 2015). Here, we have shown that in an outcrossing *A. alpina* population, genes expressed at the gametophytic stage are under stronger purifying selection than those expressed in the sporophyte. This pattern does not appear to be mainly driven by haploid selection, as individual data sets for haploid components of the gametophyte showed contrasting signatures of selection, with no consistent evidence for more efficient selection than for genes expressed in diploid tissues. This result also does not seem to be driven by antagonistic selection pressures in haploid and diploid stages, as it held for gene sets exclusively expressed in the haploid or diploid state.

By contrasting gene sets expressed in individual haploid gametophyte components, we have shown that vegetative pollen-expressed genes experienced more efficient purifying selection in comparison to genes expressed in pollen tubes and sperm cells (fig. 3). This result held after controlling for differences in expression level and breadth (fig. 4), suggesting that the contrast between vegetative pollen cells and sperm is not primarily determined by expression-related variables. A previous study by Arunkumar et al. (2013) invoked the joint contribution of haploid purging and gametophytic competition to explain the higher efficacy of selection acting on pollen-exclusive genes than on sporophyte-exclusive genes in *Capsella grandiflora*. Additionally, Harrison et al. (2019) suggested that purifying selection is currently less effective in pollen-specific than in sporophyte-specific genes in *Arabidopsis thaliana*, primarily because relaxed selection on tissue-specific genes outweighs other selective pressures in this highly self-fertilizing species.

While estimates of the DFE varied across populations in our study, purifying selection was consistently more efficient on vegetative pollen-expressed than on sperm-expressed genes across our study populations, and these two were more selectively constrained than genome-wide estimates (supplementary fig. 13). These findings agree with the suggestion that genes with a reproductive function are under strong selective constraint (Dean et al. 2009; Arunkumar et al. 2013; Finseth et al. 2014). The results further lend support to the idea that vegetative pollen and sperm cells face different selective regimes due to their contrasting roles during fertilization (Arunkumar et al. 2013; Gossmann et al. 2014). This is in line with experimental work that showed that sperm cells can be experimentally removed from pollen grains without interfering with pollen tube growth (Zhang et al. 2017), and that polyspermy avoidance mechanisms limit the possibility for competition among sperm cells (reviewed by Dresselhaus and Franklin-Tong 2013).

Last, it is important to notice that although we relied on publicly available expression data from *A. thaliana* to characterize the transcriptome of the different cell types/tissues, our analyses suggest that we have reliably inferred patterns of purifying selection differentially impacting some of the categories of genes central to our study (i.e. pollen and pistils). On the other hand, our estimates of positive selection showed some sensitivity to the source of expression data, which could partly be an effect of sampling, but still suggests that these estimates should be cautiously interpreted.

### Gene Expression Variables are not Primarily Driving the Pattern of Contrasting Selection Between Vegetative Pollen and Sperm

Contrasting signatures of selection on vegetative pollen and sperm-expressed genes have previously been reported both in outcrossing (Arunkumar et al. 2013) and selfing species (Gossmann et al. 2014). Sexual selection was invoked in both studies as the most likely explanation for this result, but expression specificity and level were proposed as important contributors to these differences. More specifically, Arunkumar et al. (2013) predicted that signatures of relaxation in sperm-expressed genes could result from their lower expression levels, because broadly and/or highly expressed genes tend to be under stronger purifying selection (Wright et al. 2004; Slotte et al. 2011; Yang and Gaut 2011; Zhang and Yang 2015). Here, we have shown that contrasting signatures of purifying selection between vegetative pollen and sperm remain after controlling for expression level and specificity (fig. 4), which confirms that the pattern is not solely driven by expression disparities among cell types. However, there were more exclusively expressed genes in sperm than in vegetative pollen (8.08% vs 3.47%, respectively), and in line with previous studies (Slotte et al. 2011; Arunkumar et al. 2013), we estimated a lower efficacy of purifying selection on exclusively expressed genes (0 < *N_e_s* < 1: all sperm-expressed genes=16.33±0.19%, exclusively sperm-expressed genes=19.46±0.20).

Taken together, our results suggest that the reduced involvement of sperm cells in pollen competition and a higher proportion of exclusively expressed genes could jointly contribute to weaker purifying selection on sperm-expressed genes.

### Low Polymorphism and Inbreeding are Associated with Weaker Purifying Selection on Male Gametophyte-Expressed Genes

Mating system variation and demographic changes are important determinants of *N_e_* in *A. alpina* populations (Laenen et al. 2018), and one or several transitions from outcrossing to mixed mating and predominant selfing have occurred in Central European and Scandinavian populations (Ansell et al. 2008; Tedder et al. 2011; Toräng et al. 2017; Laenen et al. 2018). The prediction that pollen reproductive performance will decrease in selfing lineages in comparison to their outcrossing ancestors (Mazer et al. 2010) agrees with theoretical models of density-dependence in sexual selection according to which *N_e_* drives the strength of intrasexual conflict (Kokko and Rankin 2006). In agreement with this expectation, we showed that less polymorphic and more inbred populations of *A. alpina*, which are likely to undergo higher levels of self-fertilization, experience weaker purifying selection at genes expressed in male gametophyte components than predominantly outcrossing populations (fig. 5). Importantly, this association was weaker or absent genome-wide. This suggests that weaker purifying selection on gametophytic components in more inbred populations is not solely an effect of a genome wide relaxed purifying selection in inbred populations (fig. 5, table 1). Collectively, our findings are consistent with the expectation that the intensity of sexual selection during fertilization decreases with reduced genetic diversity and a reduced number of genetically different competing haplotypes in predominantly self-fertilizing populations. Our findings also agree with a recent study on *A. thaliana* genomic data which inferred that after *A. thaliana* shifted from outcrossing to selfing 1 million years ago, genes expressed in pollen have accumulated a higher proportion of slightly deleterious mutations in comparison to sporophytic genes (Harrison et al. 2019).

### Contrasting Signatures of Sexual Selection among Male Gametophyte Components

Pollen tube competition is commonly described as the most intense stage of male-male competition (Bernasconi et al. 2004; Lankinen and Karlsson Green 2015), but this is not reflected in our results. Indeed, in our analyses, pollen tube-expressed genes were consistently inferred to experience intermediate purifying selection pressures relative to genes expressed in vegetative pollen and sperm cells, although differences were relatively subtle. Interestingly, *in vitro* experiments with *A. thaliana* indicate that the percentage of pollen germinating steadily increases from 20 to 75% during the first six hours after pollination (Fan et al. 2001), suggesting that this could be the stage at which sexual selection is likely to act by removing alleles that impair the capacity of pollen to hydrate and germinate. Additionally, it has been suggested that pollen grains from the same cohort present intrinsic differences in their capacity to hydrate and germinate, causing an asynchrony in the start for pollen tube growth (Thomson 1989). This would imply that the early stages of competition could be more intense than pollen tube growth itself, especially if the intensity of selection is density-dependent. Our results are complementary to previous analyses in *A. thaliana* in suggesting that vegetative pollen cells and the pollen tubes derived from those cells indeed play different roles during prezygotic competition, with vegetative pollen cell-expressed genes being more heavily involved in the first steps of the race for fertilization (Pina et al. 2005, Borges et al. 2008).

In this study, after controlling for transcript ubiquity and abundance, we found that patterns of positive selection differed significantly between vegetative pollen- and pollen tube-expressed genes (fig. 4c-d). This is in contrast with similar analyses conducted on *A. thaliana* suggesting that pollen- and pollen tube-expressed genes show very similar signatures of selection (Gossmann et al. 2014).

One limitation of our study, shared with previously published results (e.g. Arunkumar et al. 2013), is that we are using gene expression data from *A. thaliana*, assuming a similar expression profile in *A. alpina*. While recent comparative transcriptomic analyses suggest broad conservation of organ-specific transcriptomes in plants (Julca et al. 2021), previous studies have suggested that pollen tube transcriptomes can be species-specific (Gossmann et al. 2014; Leydon et al. 2017). This limitation could have affected our capacity to adequately identify genes that are relevant to pollen tube functioning in our focal species, and therefore, we cannot exclude the possibility that we have underestimated positive selection on pollen tube-expressed genes. Additionally, because duplicated genes can experience selective regimes than non-duplicated genes, which could confound our comparisons of gene sets (reviewed by Innan and Kondrashov 2010), we excluded paralogs from our analyses. This approach might have removed neo-functionalized genes from our data sets, which in animals have evolved reproductively relevant functions (e.g. Dai et al. 2008; Ding et al. 2010). Because there is great potential for sexual selection to operate on pollen-tube expressed genes that regulate gamete-gamete recognition and interactions with the transmitting tissues of the pistil (Leydon et al. 2017; Johnson et al. 2019; Tonnabel et al. 2021), future work would benefit from characterizing and estimating the strength of selection on species-specific pollen tube transcriptomes. Moreover, in contrast to our results on purifying selection, adaptive substitution rate estimates seem to be more sensitive to the source used to identify genes expressed on a given cell/tissue (e.g. pollen and pistils) (supplementary fig. 15b-c, Supplementary Material) once again highlighting the importance of further work on this topic.

Although sperm-expressed genes show significantly lower levels of purifying selection than vegetative pollen across populations (supplementary fig. 13, Supplementary Material), it is interesting that purifying selection on this set of genes covaries with genetic diversity and inbreeding levels. This might imply that some genes expressed in sperm are indeed subject to sexual selection, although it is difficult to completely rule out genome-wide effects on selection in less polymorphic and more inbred populations. However, there is experimental evidence that sperm-specific genes do intervene in pollen tube guidance (e.g. von Besser et al. 2006). In the case of the dataset we used in our analyses, functional classification studies identified a relative enrichment of the terms DNA repair, ubiquitin-mediated proteolysis and cell cycle progression (Borges et al. 2008). Interestingly, ubiquitin-mediated degradation has been linked to several cellular level response stimuli, including hormonal signaling (Sharma et al. 2016). Even if the removal of sperm cells does not affect pollen tube development (Zhang et al. 2017), our results suggest that the competitive ability of male gametophytes might be impaired in selfing populations due to the reduced capacity of selection to remove slightly deleterious mutations from sperm-expressed genes.

### Signatures of Purifying Selection on Synergid-Expressed Genes

Our work focused on the importance of sexual selection in driving the evolution of genes expressed in male gametophytes, however, there is a growing interest in the impact of sexual selection on female gametophyte evolution (reviewed by Beaudry et al. 2020; Tonnabel et al. 2021). For instance, a recent study on genomic data from four *Solanum* species with contrasting mating system reported that female-specific (style and ovule) genes evolve faster than male-specific genes (pollen) even after controlling for gene expression breadth (Moyle et al. 2020), although the contribution of relaxed purifying selection and positive selection to elevated divergence remains unclear.

Here, we found that in an outcrossing population, genes expressed in the synergid cells of the female gametophyte experienced stronger purifying selection and had a higher rate of adaptive fixations, but not higher *d_N_/d_S_* than any of the other datasets evaluated (fig. 3, supplementary table 2, Supplementary Material). Synergids are known to play a pivotal role in guiding pollen tubes towards the embryo sac (Higashiyama et al. 2001), and given the importance of their function in facilitating successful fertilization, we might expect these genes to be under strong evolutionary constraint. Indeed, contrary to the pattern observed for the male gametophyte, we found no correlation between genetic polymorphism and the fraction of nearly neutral new nonsynonymous mutations (supplementary fig. 14, Supplementary Material), possibly indicating that the essential functions of synergid cells in guiding pollen tubes impose strong and general constraints on these genes. More studies are required to better understand if female gametophyte-expressed genes are indeed subject to stronger constraint compared to those expressed in male gametophytes.

### Conclusions

Here, we studied the signatures of selection on genes involved in the functioning of male gametophytes and their components using *A. alpina* genomic data from populations with varying levels of polymorphism and inbreeding. We found evidence for more efficient purifying selection on genes expressed in the vegetative cell of pollen compared to pollen tubes, and significantly weaker selection on genes expressed in sperm cells. Importantly, these results cannot be attributed to differences in tissue specificity or expression levels. Purifying selection on male-gametophyte-expressed genes is stronger in predominantly outbred populations with higher levels of polymorphism than in predominantly selfing, less polymorphic populations. This result is consistent with expected effects of sexual selection at the pollen stage, although we cannot completely rule out a contribution of elevated self-fertilization on selection genome-wide. Interestingly, stringent levels of purifying selection acting on vegetative pollen-expressed genes persist even in inbred populations with low levels of polymorphism, suggesting that the reproductive functions of these genes result in strong selective constraints on their evolution.

## Materials and Methods

### Samples and Sequencing

We analyzed whole genome sequences of 228 individuals of *A. alpina* representing 13 different sampling localities (supplementary fig. 7a, supplementary table 3, Supplementary Material). We extracted DNA from dried leaves using the Qiagen DNeasy Plant Mini kit (Qiagen, Inc., Valencia, CA, USA). Sequencing libraries were prepared using the TruSeq Nano DNA sample preparation kit (Illumina Inc., San Diego, CA, USA) targeting an insert size of 350 bp. Sequencing of paired-end 150bp reads was performed on a HiSeqX (Illumina Inc.) machine with v2.5 sequencing chemistry. On average, we obtained 29 Gbp per sample (ranging from 12 to 66 Gbp) with Phred quality score above 30. A full description of sequence processing, mapping, variant calling and filtering is given in Supplementary Note 2.

### Estimates of Genetic Diversity and Inbreeding

We calculated nucleotide diversity (*π*) (Nei and Li 1979) in 10 Kb windows using pixy (https://pixy.readthedocs.io/en/1.0.0.beta1/index.html) (Korunes and Samuk 2021).

Tajima’s D (Tajima 1989) and the mean inbreeding coefficient (*F_IS_*) (Wright 1949) for each population was calculated using VCFtools (0.1.16) (Danecek et al. 2011), whereas the calculation of Watterson’s estimator of *θ* (Fu 1994) was conducted in R (R Core Team 2019). Each of these estimates were obtained for 0 and 4-fold degenerate sites identified using the *A. alpina* V5 reference genome and a lift over annotation (Arabis_alpina.MPIPZ.V5.chr.all.liftOverV4.v3.gff3, downloaded from http://www.arabis-alpina.org, last accessed July of 2018) (Willing et al. 2015) using the script NewAnnotateRef.py (https://github.com/fabbyrob/science/tree/master/pileup_analyzers, last accessed July of 2018) (Williamson et al. 2014). Genes that mapped to more than one region of the V5 reference genome based on the lift over annotation file were discarded from downstream analyses. The proportion of the genome in runs of homozygosity above 500 Kb (*F*_ROH_) was calculated for the entire data set of bi-allelic SNPs without missing data. Files were processed using BCFtools/RoH (Narasimhan et al. 2016), and the sum of RoH and *F*_ROH_ were further calculated in R (R Core Team 2019).

Estimates of average *π* across sets of genes for population Gre2 were obtained using pixy with the option --bed_file to indicate coordinates of gene sets (with separate input vcf files filtered to only retain 0-fold and 4-fold sites to obtain estimates of nucleotide diversity at each site class).

### Population Structure

We assessed population structure using fastSTRUCTURE (Raj et al. 2014). Briefly, we analyzed 1,187,118 variants retained after pruning all SNPs based on linkage disequilibrium estimated in 50 Kb windows, a step size of 5 Kb and r^2^ threshold of 0.5 using PLINK (Chang et al. 2015; Purcell and Chang 2020). We ran five replicates of fastSTRUCTURE for different *K* values ranging from 2 to 20, and for each replicate, we chose the number of populations based on the number of relevant components required to explain the structure in the dataset, as proposed by Raj et al. (2014). Ancestry results were plotted using the R package pophelper (Francis 2017) to depict the number of *K* most commonly inferred among the five replicates we ran.

### Characterization of Male and Female Gametophyte Transcriptomes

All analyzed lists of genes expressed in each tissue or cellular type were identified based on transcriptomic data from *Arabidopsis thaliana* produced with the A-AFFY-2 Affymetrix ATH1 array (ATH1-121501) obtained from publicly available datasets of the ArrayExpress repository of EMBL EBI (http://www.ebi.ac.uk/arrayexpress/) (supplementary table 5, Supplementary Material) (Borges et al. 2008; Qin et al. 2009; Wuest et al. 2010; Schmid et al. 2012). The set of genes expressed in the male gametophyte includes those present in vegetative pollen, pollen tubes and sperm cells, whereas that of the female function is based on the transcriptome of synergids, eggs and nucellus. To identify genes expressed in the sporophyte, we identified all genes called as present in the transcriptomes of vegetative shoot apex, leaf, petiole and root, and randomly selected 10,000 of them to conduct our analyses (supplementary table 6, Supplementary Material).

To convert GeneChip probe level data into expression data we used the function mas5calls.AffyBatch implemented in the R package affy (Gautier et al. 2004) that performs background correction, normalizes the values per individual array and detects probes expressed on each array based on a Wilcoxon signed rank-based algorithm. Genes were called as present with a cutoff value of *P*<0.01. We used the annotation of the ATH1-121501 probe array (https://www.ncbi.nlm.nih.gov/geo/, last updated in June 2017) to associate each element of the array with a unique *Arabidopsis* Genome Initiative (AGI) ID corresponding to the gene. The orthologs in *Arabis alpina* were identified using the one-to-one orthologs list with *A. thaliana* (Aa_At.1×1_orthologs.txt, downloaded from http://www.arabis-alpina.org/refseq.html, last accessed July of 2018) (Willing et al. 2015), and we obtained the coordinates of each gene using the lift-over annotation. Gene codes associated with two or more different coordinates (e.g. paralogs) were excluded from further analyses.

### Estimates of Purifying and Positive Selection

We inferred the distribution of negative fitness effects (DFE) of new mutations and the rate of adaptive molecular evolution (*α* and *ω_a_*) using the methods implemented in DFE-alpha v.2.15 with the programs est_dfe (Keightley and Eyre-Walker 2007) and est_alpha_omega (Eyre-Walker and Keightley 2009) respectively. Non-synonymous 0-fold degenerate sites were considered to be under selection, whereas synonymous 4-fold degenerate sites were assumed to evolve neutrally.

For analyses of population Gre2, we first produced folded rescaled SFS for 0-fold and 4-fold degenerate sites for each set of genes using SoFoS (https://github.com/CartwrightLab/SoFoS, last accessed February, 2020) to run est_dfe. To correct for missing data, SoFoS requires the prior parameters α and β to infer the shape of the beta distribution of the SFS. These were estimated based on sites without missing data. Data were rescaled to the maximum number of expected chromosomes for population Gre2 (2n=40). For comparisons across all populations, we obtained folded SFS for 0-fold and 4-fold degenerate sites without missing data, without SoFoS rescaling.

Demographic dynamics are expected to shape the SFS of both selected and neutrally evolving sites, and est_dfe implements a simple population size change model to correct for this. All analyses were done both assuming a constant population and with a two-epoch model including a population size change. In both cases, the mean effect of deleterious mutations (E(s)) and the shape parameter of the gamma distribution that describes DFE (β) were set to variable in likelihood maximization. As for analyses conducted under the two-epoch model, we estimated the population size in the second epoch (n_2_) and the duration of epoch after first population size change (t_2_). The starting values of n_2_, t_2_, E(s)) and β were set to random within the intervals specified in est_dfe instructions, and after running five independent replicates, the result with the highest likelihood was kept. Finally, we selected the best demographic model for this population using the Akaike information criterion (AIC). Based on the results obtained with est_dfe, new mutations were classified in three categories of the product of the effective population size, *N_e_*, and the strength of selection, *s*, (*N_e_s*): nearly neutral (0 < *N_e_s* < 1), moderately (1 < *N_e_s* < 10), and strongly deleterious (*N_e_s* > 10).

To estimate the proportion of adaptive substitutions (*α*) and the relative rate of adaptive substitution (*ω_α_*), we first assessed divergence using whole genome alignments between *A. alpina* and the two closely related species as outgroups: the annual *A. montbretiana* and the perennial *A. nordmanniana* (Karl and Koch 2013; Kiefer et al. 2017). Autonomous seed set in the greenhouse indicating self-compatibility has been reported for *A. montbretiana* (Kiefer et al. 2017), but no information on outcrossing levels in the field is currently available for the outgroups. We downloaded draft assemblies of *A. montbretiana* and *A. nordmanniana* from GeneBank (https://www.ncbi.nlm.nih.gov/genome) under the accession codes GCA_001484125.1 and GCA_001484925.1 respectively, and masked them as explained before for the reference genome of *A. alpina*. Pairwise alignment between genomes was executed with the lastz wrapper function implemented in the R package CNEr (Tan et al. 2019). The resulting alignments were joined with MULTIZ (Blanchette 2004) and refined using ClustalW (Thompson et al. 1994). To aid the polarization of alleles at each locus, we only kept loci with the same allele across these three species (i.e., used for the counts of total nonsynonymous and synonymous sites) and those equal in the two outgroups but different in *A. alpina*. Loci in the latter category were used to assign sites as ancestral or derived and thus count for the number of differences for 0 and 4-fold degenerate sites for each evaluated set of genes. For each gene set included in this study, we calculated DFE-alpha for the entire set of genes, and for 200 bootstrap replicates obtained by resampling over genes. To test for significant differences in the DFE, *α* and *ω_α_* between sets of genes, we performed Kruskal-Wallis tests and Dunn’s test of multiple comparisons using rank sums, with two-sided P-values adjusted using the Bonferroni method as implemented in the R package FSA (Ogle et al. 2020).

### Selection on Male Gametophyte Components and Assessment of Haploid Selection

Gene sets expressed in male and female gametophytes were split into their components to conduct DFE-alpha analyses on each of them independently. We compared these results with estimates derived from genes expressed in diploid nucellus (i.e. the tissue surrounding the ovule) (Schmid et al. 2012) and diploid unpollinated pistils (Boavida et al. 2011) (supplementary table 5, Supplementary Material) to investigate the impact of ploidy on DFE and positive selection.

### Controlling for Gene Expression Breadth and Transcript Abundance

To control for expression breadth, we intersected gene sets of the vegetative pollen cell, pollen tubes and sperm with the set of representative sporophytic tissues (a random selection of 10,000 genes vegetative shoot apex, leaf, petiole and root, supplementary table 6, Supplementary Material). This resulted in a set of genes exclusive to each component (fig. 2b). Likewise, to make this analysis more comparable to previous findings by Arunkumar et al. (2013), we produced lists of exclusively expressed genes after intersecting each set of genes expressed in the components of male gametophytes with a set of genes expressed in three-week-old seedlings (ArrayExpress dataset E-ATMX-35, Borges et al. 2008). We used a matched group comparison to control for the effect of transcript abundance. Specifically, as in Steige et al. (2017) we relied on expression data for 16,376 genes across 13 different tissues from *A. thaliana* (Schmid et al. 2005; Borges et al. 2008; Qin et al. 2009; Wuest et al. 2010; Schmid et al. 2012) (supplementary table 6, Supplementary Material). We used the maximum expression level estimated using the Robust Multi-array Average (RMA) procedure with the function expresso from the R package affy (Gautier et al. 2004) and subsampled the original gene sets were to obtain gene sets that match the distribution of maximum expression values in the set with the lowest number of genes (i.e. sperm cells) (supplementary fig. 6, Supplementary Material). DFE-alpha analyses were rerun as described above on these reduced gene sets to assess the impact of gene expression breadth and level on our estimates.

### Estimates of Purifying Selection Acting on Male Gametophytes and Synergids Across Populations

To study whether genetic polymorphism and inbreeding impact the efficiency of selection acting on genes expressed in vegetative pollen, pollen tubes and sperm, we estimated the proportion of nearly neutral new mutations (0 < *N_e_s* < 1) at 0-fold nonsynoymous sites for each gene set and across populations with contrasting estimates of *π*, *F_IS_* and *F*_ROH_. To assess genome-wide estimates of purifying selection, we repeated this analysis on a set of 10,000 randomly-selected genes from the genome annotation. For the analysis of all populations, we generated folded SFS of 0-fold and 4-fold degenerate sites for each list of genes based on sites without missing information. These analyses were conducted under the demographic scenarios of constant population size and a two-epoch model, and we selected the most likely model for each population using the AIC estimator based on analyses of randomly selected sporophytic genes. These analyses were conducted as described in the section *Estimates of Purifying and Positive Selection*.

### Correlation Between Genetic Polymorphism and Purifying Selection

We calculated Spearman’s rank correlation coefficient (*ρ*) between 4-fold polymorphism (*π*) and the fraction of nearly neutral sites in sets of genes expressed in vegetative pollen, pollen tubes, sperm cells, synergids and genome-wide estimates (10,000 randomly selected genes) across populations. To account for the phylogenetic non-independence of populations treated as data points in the correlation, we applied the phylogenetically independent contrasts (PIC) method (Felsenstein 1985) using the function pic available in the R package ape (Paradis et al. 2004). Both variables, 4-fold *π* and the fraction of sites in 0 < *N_e_s* < 1, were corrected using a maximum likelihood (ML) tree based on 10 *A. alpina* populations (supplementary figure 12a, Supplementary Material). We obtained this ML tree including 169 samples from ten populations using the pipeline SNPhylo (Lee et al. 2014) which reconstructs a ML tree using DNAML from the PHYLIP package (Felsenstein 1989), under the F84 model of sequence evolution (Kishino and Hasegawa 1989). SNPs were pruned based on a LD threshold of 0.5 estimated along 500 Kb sliding windows, which resulted in a set of 15,033 markers. We estimated node support based on 500 bootstrap replicates in PhyML (Guindon et al. 2010). Finally, we reduced the tree to keep a single sample per population using the function keep.tip from the ape package (Paradis et al. 2004) (supplementary figure 12b, Supplementary Material), such that each tip could be associated with the corresponding estimates of 4-fold *π* and 0 < *N_e_s* < 1 for each set of genes. Finally, we repeated the correlations analysis using the corrected values.

### Comparison of Differences in Purifying Selection Between Inbred and Outbred Populations

With the exception of one population with intermediate value for *F*_ROH_, our populations fell into two clearly separated groups in terms of *F_IS_* and mean *F*_ROH_ (mean 4-fold *F_IS_* < 0.5 and mean *F*_ROH_ < 0.5 vs. mean 4-fold *F_IS_* > 0.5 and mean *F*_ROH_ > 0.5; supplementary figures 2 and 3, Supplementary Material). Those with mean 4-fold *F_IS_* > 0.5 and mean *F*_ROH_ > 0.5 all had a mean 4-fold *F_IS_* > 0.85, which corresponds to a mean effective equilibrium self-fertilization rate of 0.92 (Wright 1969), whereas those in the mean 4-fold *F_IS_* < 0.5 and mean *F*_ROH_ < 0.5 group all had a mean 4-fold *F_IS_* < 0.04, corresponding to a mean equilibrium effective selfing rate of 0.08. We therefore tentatively classified these groups as predominantly self-fertilizing and predominantly outcrossing. Then, we tested for differences in the fraction of nearly neutral 0-fold non-synonymous mutations (0 < *N_e_s* < 1) between these two classes of populations for each data set using a Wilcoxon rank-sum test in R (R Core Team 2019).

## Supplementary Material

Notes 1-2, Tables 1-6, Figures 1-15

## Acknowledgments

We thank Verena E. Kutschera from the Swedish Bioinformatics Advisory Program (NBIS, Sweden) for advice on bioinformatic and genetic analyses, Tanja Pyhäjärvi for discussion of the first version of the manuscript, and Rhonda Snook for interesting discussion of sexual selection in plants and animals. This work was supported by the Science for Life Laboratory, Swedish Biodiversity Program. The Swedish Biodiversity Program has been made available by support from the Knut and Alice Wallenberg foundation. The computations were enabled by resources provided by the Swedish National Infrastructure for Computing (SNIC) at UPPMAX partially funded by the Swedish Research Council through grant agreement no. 2018-05973. Sequencing was performed by the SNP&SEQ Technology Platform in Uppsala. The facility is part of the National Genomics Infrastructure (NGI) Sweden and Science for Life Laboratory. The SNP&SEQ Platform is also supported by the Swedish Research Council and the Knut and Alice Wallenberg Foundation.

## Notes

### Competing Interest Statement

The authors have declared no competing interest.

## References

Ansell SW, Grundmann M, Russell SJ, Schneider H, Vogel JC. 2008. Genetic Discontinuity, Breeding-System Change and Population History of *Arabis alpina* in the Italian Peninsula and Adjacent Alps. Mol Ecol. 17:2245–2257.

Arunkumar R, Josephs EB, Williamson RJ, Wright SI. 2013. Pollen-Specific, but not Sperm-Specific, Genes Show Stronger Purifying Selection and Higher Rates of Positive Selection than Sporophytic Genes in *Capsella grandiflora*. Mol Biol Evol. 30:2475–2486.

Austerlitz F, Gleiser G, Teixeira S, Bernasconi G. 2012. The Effects of Inbreeding, Genetic Dissimilarity and Phenotype on Male Reproductive Success in a Dioecious Plant. Proc Biol Sci. 279:91–100.

Barrett SCH. 2002. The Evolution of Plant Sexual Diversity. Nature Rev Genet. 3:274–284.

Beaudry FEG, Rifkin JL, Barrett SCH, Wright SI. 2020. Evolutionary Genomics of Plant Gametophytic Selection. Plant Comm. 1:100115.

Bernasconi G, Ashman T-L, Birkhead TR, Bishop JDD, Grossniklaus U, Kubli E, Marshall DL, Schmid B, Skogsmyr I, Snook RR, et al. 2004. Evolutionary Ecology of the Prezygotic Stage. Science. 303:971–975.

Blanchette M. 2004. Aligning Multiple Genomic Sequences with the Threaded Blockset Aligner. Genome Res. 14:708–715.

Borges F, Gomes G, Gardner R, Moreno N, McCormick S, Feijó JA, Becker JD. 2008. Comparative Transcriptomics of *Arabidopsis* Sperm Cells. Plant Physiol. 148:1168–1181.

Chang CC, Chow CC, Tellier LC, Vattikuti S, Purcell SM, Lee JJ. 2015. Second-Generation PLINK: Rising to the Challenge of Larger and Richer Datasets. Gigascience. 4:7.

Charlesworth D, Charlesworth B. 1992. The Effects of Selection in the Gametophyte Stage on Mutational Load. Evolution. 46:703–720.

Cutter AD. 2019. Reproductive Transitions in Plants and Animals: Selfing Syndrome, Sexual Selection and Speciation. New Phytol. 224:1080–1094.

Clark NL, Aagaard JE, Swanson WJ. 2006. Evolution of Reproductive Proteins from Animals and Plants. Reproduction. 131:11–22.

Dai H, Chen Y, Chen S, Mao Q, Kennedy D, Landback P, Eyre-Walker A, Du W, Long M. 2008. The Evolution of Courtship Behaviors Through the Origination of a New Gene in *Drosophila*. Proc Natl Acad Sci U S A. 105:7478–83.

Danecek P, Auton A, Abecasis G, Albers CA, Banks E, DePristo MA, Handsaker RE, Lunter G, Marth GT, Sherry ST, et al. 2011. The Variant Call Format and vcftools. Bioinformatics. 27:2156–2158.

Dapper AL, Wade MJ. 2016. The Evolution of Sperm Competition Genes: the Effect of Mating System on Levels of Genetic Variation Within and Between Species. Evolution. 70:502–11.

Dapper AL, Wade MJ. 2020. Relaxed Selection and the Rapid Evolution of Reproductive Genes. Trends Genet. 36:640–649.

Darwin C. 1871. The Descent of Man, and Selection in Relation to Sex. London: John Murray.

Darwin C. 1876. The Effects of Cross and Self Fertilisation in the Vegetable Kingdom. London: John Murray.

Dean MD, Clark NL, Findlay GD, Karn RC, Yi X, Swanson WJ, MacCoss MJ, Nachman MW. 2009. Proteomics and Comparative Genomic Investigations Reveal Heterogeneity in Evolutionary Rate of Male Reproductive Proteins in Mice *(Mus domesticus)*. Mol Biol Evol. 26:1733–1743.

Ding Y, Zhao L, Yang S, Jiang Y, Chen Y, Zhao R, Zhang Y, Zhang G, Dong Y, Yu H, et al. 2010. A Young *Drosophila* Duplicate Gene Plays Essential Roles in Spermatogenesis by Regulating Several Y-Linked Male Fertility Genes. PLOS Genet. 6:e1001255.

Dresselhaus T, Franklin-Tong N. 2013. Male–Female Crosstalk during Pollen Germination, Tube Growth and Guidance, and Double Fertilization. Mol Plant. 6:1018–1036.

Dresselhaus T, Sprunck S, Wessel GM. 2016. Fertilization Mechanisms in Flowering Plants. Curr Biol. 26:R125–R139.

Drummond DA, Bloom JD, Adami C, Wilke CO, Arnold FH. 2005. Why Highly Expressed Proteins Evolve Slowly. Proc Natl Acad Sci U S A. 102:14338–14343.

Eyre-Walker A, Keightley PD. 2009. Estimating the Rate of Adaptive Molecular Evolution in the Presence of Slightly Deleterious Mutations and Population Size Change. Mol Biol Evol. 26:2097–2108.

Fan L-M, Wang Y-F, Wang H, Wu W-H. 2001. *In vitro Arabidopsis* Pollen Germination and Characterization of the Inward Potassium Currents in *Arabidopsis* Pollen Grain Protoplasts. J Exp Bot. 52:1603–1614.

Felsenstein J. 1985. Phylogenies and the Comparative Method. Am Nat. 125:1–15.

Felsenstein J. 1989. PHYLIP-Phylogeny Inference Package (Ver. 3.2). Cladistics. 5:164–166.

Finseth FR, Bondra E, Harrison RG. 2014. Selective Constraint Dominates the Evolution of Genes Expressed in a Novel Reproductive Gland. Mol Biol Evol. 31:3266–3281.

Francis RM. 2017. pophelper: an R Package and Web App to Analyse and Visualize Population Structure. Mol Ecol Resour. 17:27–32.

Fu YX. 1994. Estimating Effective Population Size or Mutation Rate Using the Frequencies of Mutations of Various Classes in a Sample of DNA Sequences. Genetics. 138:1375–1386.

Gautier L, Cope L, Bolstad BM, Irizarry RA. 2004. affy-Analysis of Affymetrix GeneChip Data at the Probe Level. Bioinformatics. 20:307–315.

Gerstein AC, Otto SP. 2009. Ploidy and the Causes of Genomic Evolution. J Hered. 100:571–581.

Gossmann TI, Schmid MW, Grossniklaus U, Schmid KJ. 2014. Selection-Driven Evolution of Sex-Biased Genes is Consistent with Sexual Selection in *Arabidopsis thaliana*. Mol Biol Evol. 31:574–583.

Guindon S, Dufayard JF, Lefort V, Anisimova M, Hordijk W, Gascuel O. 2010. New Algorithms and Methods to Estimate Maximum-Likelihood Phylogenies: Assessing the Performance of PhyML 3.0. Syst Biol. 59:307–21.

Haldane JBS. 1932. The Causes of Evolution. Princeton, NJ: Princeton University Press.

Haldane JBS. 1933. The Part Played by Recurrent Mutation in Evolution. Am Nat. 67:5–19.

Harrison MC, Mallon EB, Twell D, Hammond RL. 2019. Deleterious Mutation Accumulation in *Arabidopsis thaliana* Pollen Genes: a Role for a Recent Relaxation of Selection. Genome Biol Evol. 11:1939–1951.

Hartfield M, Bataillon T, Glémin S. 2017. The Evolutionary Interplay Between Adaptation and Self-Fertilization. Trends Genet. 33:420–431.

Higashiyama T, Yabe S, Sasaki N, Nishimura Y, Miyagishima S, Kuroiwa H, Kuroiwa T. 2001. Pollen Tube Attraction by the Synergid Cell. Science. 293:1480–1483.

Honys D, Twell D. 2004. Transcriptome Analysis of Haploid Male Gametophyte Development in *Arabidopsis*. Genome Biol. 5:R85.

Hove AA, Mazer SJ. 2013. Pollen Performance in *Clarkia* Taxa with Contrasting Mating Systems: Implications for Male Gametophytic Evolution in Selfers and Outcrossers. Plants. 2:248–278.

Immler S, Arnqvist G, Otto SP. 2012. Ploidally Antagonistic Selection Maintains Stable Genetic Polymorphism. Evolution. 66:55–65.

Innan H, Kondrashov F. 2010. The Evolution of Gene Duplications: Classifying and Distinguishing Between Models. Nat Rev Genet. 11:97–108.

Johnson MA, Harper JF, Palanivelu R. 2019. A Fruitful Journey: Pollen Tube Navigation from Germination to Fertilization. Annu Rev Plant Biol. 29:809–837.

Julca I, Ferrari C, Flores-Tornero M, Proost S, Lindner AC, Hackenberg D, Steinbachová L, Michaelidis C, Gomes Pereira S, Misra CS, et al. 2021. Comparative Transcriptomic Analysis Reveals Conserved Programmes Underpinning Organogenesis and Reproduction in Land Plants. Nat Plants. 7:1143–1159.

Karl R, Koch MA. 2013. A World-Wide Perspective on Crucifer Speciation and Evolution: Phylogenetics, Biogeography snd Trait Evolution in Tribe Arabideae. Ann Bot. 112:983–1001.

Kasahara RD, Maruyama D, Hamamura Y, Sakakibara T, Twell D, Higashiyama T. 2012. Fertilization Recovery After Defective Sperm Cell Release in *Arabidopsis*. Curr Biol. 22:1084–1089.

Keightley PD, Eyre-Walker A. 2007. Joint Inference of the Distribution of Fitness Effects of Deleterious Mutations and Population Demography Based on Nucleotide Polymorphism Frequencies. Genetics. 177:2251–2261.

Kiefer C, Severing E, Karl R, Bergonzi S, Koch M, Tresch A, Coupland G. 2017. Divergence of Annual and Perennial Species in the Brassicaceae and the Contribution of *cis*-Acting Variation at *FLC* Orthologues. Mol Ecol. 26:3437–3457.

Kishino H, Hasegawa M. 1989. Evaluation of the Maximum Likelihood Estimate of the Evolutionary Tree Topologies from DNA Sequence Data, and the Branching Order in Hominoidea. J Mol Evol. 29:170–179.

Kokko H, Rankin DJ. 2006. Lonely Hearts or Sex in the City? Density-Dependent Effects in Mating Systems. Philos Trans R Soc Lond Ser B Biol Sci. 361:319.

Kondrashov AS, Crow JF. 1991. Haploidy or Diploidy: Which is Better? Nature. 351:314–315.

Korunes KL and Samuk K. (2021). pixy: Unbiased estimation of nucleotide diversity and divergence in the presence of missing data. Mol Ecol Resour. doi: 10.1111/1755-0998.13326. Online ahead of print.

Laenen B, Tedder A, Nowak MD, Toräng P, Wunder J, Wötzel S, Steige KA, Kourmpetis Y, Odong T, Drouzas AD, et al. 2018. Demography and Mating System Shape the Genome-Wide Impact of Purifying Selection in *Arabis alpina*. Proc Natl Acad Sci U S A. 115:816–821.

Lankinen Å, Hydbom S, Strandh M. 2017. Sexually Antagonistic Evolution Caused by Male-Male Competition in the Pistil. Evolution. 71: 2359–2369.

Lankinen Å, Karlsson Green K. 2015. Using Theories of Sexual Selection and Sexual Conflict to Improve our Understanding of Plant Ecology and Evolution. AoB Plants. 7:plv008.

Lankinen Å, Skogsmyr I. 2002. Pollen Competitive Ability: the Effect of Proportion in Two-Donor Crosses. Evol Ecol Res. 4:687–700.

Lee TH, Guo H, Wang X, Kim C, Paterson AH. 2014. SNPhylo: a Pipeline to Construct a Phylogenetic Tree from Huge SNP Data. BMC Genomics. 15:162.

Leydon AR, Weinreb C, Venable E, Reinders A, Ward JM, Johnson MA. 2017. The Molecular Dialog between Flowering Plant Reproductive Partners Defined by SNP-Informed RNA-Sequencing. Plant Cell. 29:984–1006.

Lohani N, Singh MB, Bhalla PL. 2021. RNA-Seq Highlights Molecular Events Associated with Impaired Pollen-Pistil Interactions Following Short-Term Heat Stress in *Brassica napus*. Front Plant Sci. 11:622748.

Mattila TM, Laenen B, Horvath R, Hämälä T, Savolainen O, Slotte T. 2019. Impact of Demography on Linked Selection in Two Outcrossing Brassicaceae Species. Ecol Evol. 9:9532–9545.

Mazer SJ, Hove AA, Miller BS, Barbet-Massin M. 2010. The Joint Evolution of Mating System and Pollen Performance: Predictions Regarding Male Gametophytic Evolution in Selfers vs. Outcrossers. Perspect Plant Ecol Evol Syst. 12:31–41.

Mazer SJ, Hendrickson BT, Chellew JP, Kim LJ, Liu JW, Shu J, Sharma MV. 2018. Divergence in Pollen Performance Between *Clarkia* Sister Species with Contrasting Mating Systems Supports Predictions of Sexual Selection. Evolution. 72:453–472.

Moyle LC, Wu M, Gibson MJS. 2021. Reproductive Proteins Evolve Faster than Non-Reproductive Proteins Among *Solanum* Species. Front Plant Sci. 12:635990.

Narasimhan V, Danecek P, Scally A, Xue Y, Tyler-Smith C, Durbin R. 2016. BCFtools/RoH: a Hidden Markov Model Approach for Detecting Autozygosity from Next-Generation Sequencing Data. Bioinformatics. 32:1749–1751.

Nei M, Li WH. 1979. Mathematical Model for Studying Genetic Variation in Terms of Restriction Endonucleases. Proc Natl Acad Sci U S A. 76:5269–5273.

Ogle DH, Wheeler P, Dinno A. 2020. FSA: Fisheries Stock Analysis. https://github.com/droglenc/FSA.

Palanivelu R, Tsukamoto T. 2012. Pathfinding in Angiosperm Reproduction: Pollen Tube Guidance by Pistils Ensures Successful Double Fertilization. Wiley Interdiscip Rev Dev Biol. 1:96–113.

Pannell JR and Labouche A. 2013. The Incidence and Selection of Multiple Mating in Plants. Phil Trans R Soc B. 368: 20120051.

Paradis E, Claude J, Strimmer K. 2004. APE: Analyses of Phylogenetics and Evolution in R language. Bioinformatics. 20:289–290.

Pasonen HL, Pulkkinen P, Kapyla M, Blom A. 1999. Pollen-Tube Growth Rate and Seed-Siring Success Among *Betula pendula* Clones. New Phytol. 143:243–251.

Peters MAE, Weis AE. 2018. Selection for Pollen Competitive Ability in Mixed-Mating Systems. Evolution. 72:2513–2536.

Petrén H, Toräng P, Ågren J, Friberg M. 2021. Evolution of Floral Scent in Relation to Self-Incompatibility and Capacity for Autonomous Self-Pollination in the Perennial Herb *Arabis alpina*. Ann Bot. 127:737–747.

Pina C, Pinto F, Feijó JA, Becker JD. 2005. Gene Family Analysis of the *Arabidopsis* Pollen Transcriptome Reveals Biological Implications for Cell Growth, Division Control, and Gene Expression Regulation. Plant Physiol. 138:744–756.

Purcell S, Chang C. 2020. PLINK. www.cog-genomics.org/plink/2.0/

Qin Y, Leydon AR, Manziello A, Pandey R, Mount D, Denic S, Vasic B, Johnson MA, Palanivelu R. 2009. Penetration of the Stigma and Style Elicits a Novel Transcriptome in Pollen Tubes, Pointing to Genes Critical for Growth in a Pistil. PLoS Genet. 5:e1000621.

R Core Team. 2019. R: A language and environment for statistical computing. R Foundation for Statistical Computing. Vienna, Austria. URL https://www.R-project.org/.

Raj A, Stephens M, Pritchard JK. 2014. fastSTRUCTURE: Variational Inference of Population Structure in Large SNP Data Sets. Genetics. 197:573–589.

Rowe L, Houle D. 1996. The Lek Paradox and the Capture of Genetic Variance by Condition Dependent Traits. Proc R Soc Lond B. 263:1415–1421

Schmid MW, Schmidt A, Klostermeier UC, Barann M, Rosenstiel P, Grossniklaus U. 2012. A Powerful Method for Transcriptional Profiling of Specific Cell Types in Eukaryotes: Laser-Assisted Microdissection and RNA Sequencing. PLoS ONE. 7:e29685.

Sharma B, Joshi D, Yadav PK, Gupta AK, Bhatt TK. 2016. Role of Ubiquitin-Mediated Degradation System in Plant Biology. Front Plant Sci. 7:806.

Shuster SM. 2009. Sexual Selection and Mating Systems. Proc Natl Acad Sci U S A. 106:10009–10016.

Sicard A, Lenhard M. 2011. The Selfing Syndrome: A Model for Studying the Genetic and Evolutionary Basis of Morphological *Adaptation* in Plants. Ann Bot. 107:1433–1443.

Skogsmyr I, Lankinen ÅSA. 2002. Sexual Selection: an Evolutionary Force in Plants? Biol Rev. 77:537–562.

Slotte T, Bataillon T, Hansen TT, St Onge K, Wright SI, Schierup MH. 2011. Genomic Determinants of Protein Evolution and Polymorphism in *Arabidopsis*. Genome Biol Evol. 3:1210–1219.

Slotte T. 2014. The Impact of Linked Selection on Plant Genomic Variation. Brief Funct Genomics. 13:268–275.

Smith-Huerta NL. 1996. Pollen Germination and Tube Growth in Selfing and Outcrossing Populations of Clarkia tembloriensis (Onagraceae). Int J Plant Sci. 157:228–233.

Steige KA, Laenen B, Reimegård J, Scofield DG, Slotte T. 2017. Genomic analysis reveals major determinants of cis-regulatory variation in Capsella grandiflora. Proc Natl Acad Sci 114: 1087–1092.

Swanson WJ, Vacquier VD. 2002. The Rapid Evolution of Reproductive Proteins. Nat Rev Genet. 3:137–44.

Szövényi P, Ricca M, Hock Z, Shaw JA, Shimizu KK, Wagner A. 2013. Selection is No More Efficient in Haploid than in Diploid Life Stages of an Angiosperm and a Moss. Mol Biol Evol. 30:1929–1939.

Tajima F. 1989. Statistical Method for Testing the Neutral Mutation Hypothesis by DNA Polymorphism. Genetics. 123:585–595.

Tan G, Polychronopoulos D, Lenhard B. 2019. CNEr: A Toolkit for Exploring Extreme Noncoding Conservation. PLoS Comput Biol. 15:e1006940.

Taylor ML, Williams JH. 2012. Pollen Tube Development in Two Species of *Trithuria* (Hydatellaceae) with Contrasting Breeding Systems. Sex Plant Reprod. 25:83–96.

Tedder A, Ansell SW, Lao X, Vogel JC, Mable BK. 2011. Sporophytic Self-Incompatibility Genes and Mating System Variation in *Arabis alpina*. Ann Bot. 108:699–713.

Thompson JD, Higgins DG, Gibson TJ. 1994. CLUSTAL W: Improving the Sensitivity of Progressive Multiple Sequence Alignment Through Sequence Weighting, Position-Specific Gap Penalties and Weight Matrix Choice. Nucleic Acids Res. 22:4673–4680.

Thomson JD. 1989. Germination Schedules of Pollen Grains: Implications for Pollen Selection. Evolution. 43:220–223.

Tonnabel J, David P, Janicke T, Lehner A, Mollet JC, Pannell JR, Dufay M. 2021. The Scope for Postmating Sexual Selection in Plants. Trends Ecol Evol. 36:556–567.

Toräng P, Vikström L, Wunder J, Wötzel S, Coupland G, Ågren J. 2017. Evolution of the Selfing Syndrome: Anther Orientation and Herkogamy Together Determine Reproductive Assurance in a Self-Compatible Plant. Evolution. 71:2206–2218.

Varis S, Reininharju J, Santanen A, Ranta H, Pulkkinen P. 2010. Interactions During in Vitro Germination of Scots Pine Pollen. Trees. 24:99–104.

von Besser K, Frank AC, Johnson MA, Preuss D. 2006. *Arabidopsis HAP2 (GCS1)* is a Sperm-Specific Gene Required for Pollen Tube Guidance and Fertilization. Development. 133:4761–4769.

Walsh NE, Charlesworth D. 1992. Evolutionary Interpretations of Differences in Pollen Tube Growth Rates. Q Rev Biol. 67:19–37.

Willi Y. 2013. The Battle of the Sexes over Seed Size: Support for Both Kinship Genomic Imprinting and Interlocus Contest Evolution. Am Nat. 181:787–798.

Williamson RJ, Josephs EB, Platts AE, Hazzouri KM, Haudry A, Blanchette M, Wright SI. 2014. Evidence for Widespread Positive and Negative Selection in Coding and Conserved Noncoding Regions of *Capsella grandiflora*. PLoS Genet. 10:e1004622.

Willing E-M, Rawat V, Mandáková T, Maumus F, James GV, Nordström KJV, Becker C, Warthmann N, Chica C, Szarzynska B, et al. 2015. Genome Expansion of *Arabis Alpina* Linked with Retrotransposition and Reduced Symmetric DNA Methylation. Nat Plants. 1:14023.

Wright S. 1949. The Genetical Structure of Populations. Ann Eugen. 15:323–354.

Wright SI, Yau CBK, Looseley M, Meyers BC. 2004. Effects of Gene Expression on Molecular Evolution in *Arabidopsis thaliana* and *Arabidopsis lyrata*. Mol Biol Evol. 21:1719–1726.

Wright SI, Kalisz S, Slotte T. 2013. Evolutionary Consequences of Self-Fertilization in Plants. Proc R Soc B. 280:20130133

Wright S. 1969. Evolution and the Genetics of Populations, Vol 2. The Theory of Gene Frequencies. Univ. Chicago Press, Chicago, IL.

Wuest SE, Vijverberg K, Schmidt A, Weiss M, Gheyselinck J, Lohr M, Wellmer F, Rahnenführer J, von Mering C, Grossniklaus U. 2010. *Arabidopsis* Female Gametophyte Gene Expression Map Reveals Similarities Between Plant and Animal Gametes. Curr Biol. 20:506–512.

Yang L, Gaut BS. 2011. Factors that Contribute to Variation in Evolutionary Rate among *Arabidopsis* Genes. Mol Biol Evol. 28:2359–2369.

Zhang J, Huang Q, Zhong S, Bleckmann A, Huang J, Guo X, Lin Q, Gu H, Dong J, Dresselhaus T, et al. 2017. Sperm Cells are Passive Cargo of the Pollen Tube in Plant Fertilization. Nat Plants. 3:17079.

Zhang J, Yang J-R. 2015. Determinants of the Rate of Protein Sequence Evolution. Nat Rev Genet. 16:409–420.

Zheng Y-Y, Lin X-J, Liang H-M, Wang F-F, Chen L-Y. 2018. The Long Journey of Pollen Tube in the Pistil. Int J Mol Sci. 19:3529.

