## Supplementary Material for "Genomic Signatures of Sexual Selection on Pollen-Expressed Genes in *Arabis alpina*"

#### Notes

**Note 1**. Impact of the use of heterospecific expression data.
**Note 2**. Sequence Processing, Mapping, Variant Calling and Filtering

#### Tables

**Table 1.** Estimates of inbreeding and genetic diversity across populations for 0 and 4-fold degenerate sites

**Table 2.** Estimates of genetic diversity and divergence for the different gene sets for 0 and 4-fold degenerate sites

**Table 3.** Origin of 228 samples included in this study from 13 different populations

**Table 4.** Number of SNPs and invariant sites after applying each filtering step

**Table 5.** Publicly available ArrayExpress datasets used to identify genes expressed on each tissue and/or cellular type

**Table 6.** Publicly available ArrayExpress datasets used characterize the distribution of expression level for each data set

#### Figures

**Figure 1.** Map depicting mean 4-fold $\pi$ values for each population

**Figure 2.** Estimates of $F_{IS}$ per individual for 0 and 4-fold sites.

**Figure 3.** Estimates of the proportion of the genome in runs of homozygosity above 500 Kb ($F_{\mathrm{ROH}}$).

**Figure 4.** Proportion of adaptive substitutions ($\alpha$) and adaptive substitutions relative to neutral divergence ($\omega_\alpha$) between the gametophyte and sporophyte-expressed genes.

**Figure 5.** Proportion of adaptive substitutions relative to neutral divergence ($\omega_\alpha$) estimated for the components of the male and female gametophytes, exclusively expressed genes and data sets corrected for expression level.

**Figure 6.** DFE estimates, proportion of adaptive substitutions ($\alpha$) and the proportion of adaptive substitutions relative to neutral divergence ($\omega_\alpha$) between sets of genes expressed in pollen and unpollinated pistils using expression data from *Arabidopsis thaliana* and *Brassica napus* for population Gre2.

**Figure 7.** Comparison of DFE estimates for sets of genes putatively exclusive to vegetative pollen, pollen tubes, sperm cells and sporophytic tissues.

**Figure 8**. Frequency polygons depicting the distribution of expression level values for each gene set before and after fitting to the distribution of sperm-expressed genes.

**Figure 9.** Geographic origin of 13 populations of *A. alpina* included in this study and bayesian clustering of 228 individuals showing ancestral proportions based on fastSTRUCTURE results.

**Figure 10.** Correlation between $F_{\mathrm{ROH}}$, mean 4-fold $\pi$, and 4-fold $F_{IS}$ respect to the fraction of nearly neutral sites ($0 < N_e s < 1$) per data set.

**Figure 11.** Inferred DFE for 10,000 randomly selected genome-wide genes for all populations.

**Figure 12.** Correlation between PIC-corrected estimates of 4-fold $\pi$ and the fraction of nearly neutral sites ($0 < N_e s < 1$) across *A. alpina* populations.

**Figure 13.** Fraction of new nearly neutral nonsynonymous mutations, $0 < N_e s < 1$, across populations for sets of genes expressed in pollen and sperm.

**Figure 14.** Scatterplot depicting the correlation between estimates of 4-fold $\pi$ and the fraction of nearly neutral sites ($0 < N_e s < 1$) for genes expressed in the synergids.

**Figure 15.** ML-inferred phylogeny of ten populations of *A. alpina*.

**References**

### Notes

**Note 1.** Impact of the use of heterospecific expression data.

In this study we relied on gene expression data from *A. thaliana* to identify genes expressed in specific cell types and tissues, under the assumption that orthologs in *A. alpina* would show similar expression patterns (as previously done by Arunkumar et al. 2013). As no similarly detailed expression data were available for *A. alpina,* we assessed the sensitivity of our analyses to using expression data from a different Brassicaceae species. Specifically, we made use of publicly available data to estimate the degree of conservation of the transcriptomes of *A. thaliana* and *Brassica napus* for pollen and unpollinated pistil-expressed genes.

We processed a publicly available data set obtained from *Brassica napus* (Illumina RNA-Seq sequences from non-stressed control samples) (Lohani et al. 2021, supplementary Data Sheet 2, Tables S1b and S1c) to identify genes expressed in mature pollen and unpollinated pistils. This data set reported transcript abundance as Transcripts per million (TPM) for three replicates of each kind. In order to call a gene as expressed, we required an expression value equal or larger than 2 TPM (as suggested by Wagner et al. 2013) for all three replicates. To identify the orthologs of *B. napus* in *A. thaliana*, we used the list of homologs provided as supplementary material by Chalhoub et al. (2014) (supplementary Table S19), and this list was finally used to identify the corresponding *A. alpina* orthologs. All these steps were conducted in R (R Core Team 2019). We used these sets of pollen and pistil-expressed genes derived from *B. napus* and *A. thaliana* to estimate DFE, $\alpha$ and $\omega_\alpha$ using polymorphism data from *A. alpina* and divergence to two outgroups following the same procedure described in the Methods (*Estimates of Purifying and Positive Selection*).

Using *B. napus* data, we identified 2,489 pollen-expressed genes with orthologs in *A. alpina.* We found that 69.30% of these genes (1,725 genes) were also present in the set of pollen-expressed genes in *A. thaliana* (Borges et al. 2008). Likewise, the transcriptome of *B. napus* pistils (Lohani et al. 2021) was composed of 9,162 genes with an identifiable ortholog in *A. alpina.* This data set overlaps to 72.46% (6,639) with the set of genes previously classified as expressed in pistils using *A. thaliana* microarray data (Boavida et al. 2011).

We compared patterns of purifying and positive selection for each data set. Our DFE comparisons show that there are small but significant differences in the proportion of effectively neutral sites between the set of pollen-expressed genes identified using *A. thaliana* and *B. napus* data ($P$<0.001, Kruskal-Wallis test followed by Dunn's test with Bonferroni correction, based on contrasts of 200 bootstrap replicates), with 13.67±0.14% and 12.03±0.17% of new nonsynonymous mutations with $0 < N_e s < 1$ (supplementary fig. 6a, Supplementary Material). Similar results were observed for pistil-expressed genes with the set of genes identified using *A. thaliana* showing the highest proportion of effectively neutral new nonsynonymous mutations ($0 < N_e s < 1$: 15.56±0.10% and 13.97±0.09%) (supplementary fig. 6a, Supplementary Material).

As for the estimates of positive selection, we found no significant differences between the estimates of $\alpha$ for pollen-expressed genes using *A. thaliana* and *B. napus*-derived data (0.082± 0.011 and 0.083± 0.015 respectively, NS, Dunn's test based on contrasts of 200 bootstrap replicates, Bonferroni corrected $P$-values (supplementary fig. 6b, Supplementary Material). In contrast, we inferred a significantly lower proportion of adaptive substitutions in pistil-expressed genes when using *A. thaliana* than *B. napus* expression data (supplementary fig. 6b-c, Supplementary Material).

Importantly, each of the strategies used to identify pollen and pistil expressed genes led to the conclusion that purifying selection is significantly stronger in pollen-expressed than in pistil-expressed genes (0 < $N_e s$ < 1 estimates in *A. thaliana*= pollen: 13.67±0.14%, pistil: 15.56±0.10%; *B. napus*= pollen: 12.03±0.17%, pistil: 13.97±0.09%) (supplementary fig. 6a, Supplementary Material). This result shows that using *A. thaliana*-derived expression data produces results similar to those obtained using data from another Brassicaceae species (*B. napus*), suggesting that this approach is reliable and can be used to detect contrasting patterns of purifying selection that affect genes expressed in different cell types or tissues. On the other hand, estimates of positive selection can differ depending on the source used to identify pollen and pistil-expressed genes, and should be more cautiously interpreted.

**Note 2:** Sequence Processing, Mapping, Variant Calling and Filtering

Sequencing adapters were identified using cutadapt v.1.8 (Martin 2011) and trimmed from the raw sequences using Trimmomatic v.0.32 (Bolger et al. 2014). The trimmed paired-end sequence reads were mapped to the *A. alpina* V5 reference genome assembly (Jiao et al. 2017) using BWA-MEM v0.7.8 (Li and Durbin 2009). Duplicated reads were removed using MarkDuplicates in Picard tools v2.0.1 (Broad Institute 2019). Across all our samples, the mean read depth was 55X (ranging from 12 to 128X). BAM alignment files were processed using the Genome Analysis Toolkit (GATK) v.3.8.0 (McKenna et al. 2010) by restricting the analyses to the eight chromosomes. Indels were realigned using GATK RealignerTargetCreator and IndelRealigner, and SNPs were called with GATK HaplotypeCaller (Poplin et al. 2017) using the DISCOVERY genotyping mode and default parameters. Genotype calling resulted in 219,544,869 invariant positions and 34,243,876 bi-allelic SNPs.

We filtered the resulting vcf file in four successive steps as follows. First, we removed SNPs and invariant sites with more than 10% of individuals with missing data, coverage higher than 200X or lower than 8X. Second, we removed all SNPs and invariant sites present in simple and complex repetitive regions identified with RepeatMasker v.4.1.0 (Smit et al. 2019) using the database of transposable elements for *A. alpina* downloaded from RepetDB (http://urgi.versailles.inra.fr/repetdb/begin.do, last accessed May of 2019). Third, as the *A. alpina* genome is very repeat-rich (Willing et al. 2015), we conducted additional filtering to remove repetitive regions that may be collapsed in the genome assembly. Specifically, we removed SNPs and invariant sites in 500 bp windows with unusually high coverage. For each sample, read depth was calculated in 500 bp windows using the R package bamsignals v1.20.0 (Mammana and Helmuth 2020). We removed windows with a coverage 1.5 higher than the within-sample median in at least in half of all individuals. Finally, to avoid false heterozygous calls based on a low number of alternate alleles, we set as missing heterozygous genotype calls based on an allele balance ratio higher than 0.7 or lower than 0.3, and we filtered again sites with a missing data percentage higher than 10%. After these filtering steps, we retained 2,776,972 biallelic SNPs and 34,790,202 invariant positions (supplementary table 4, Supplementary Material).

### Tables

**Table 1.** Estimates of inbreeding and genetic diversity across populations for 0 and 4-fold degenerate sites.

| Population | Degeneracy | No. sites | No. SNPs | No. samples | $\pi$ | $\pi_N/\pi_S$ | Watterson's θ | Mean $F_{IS}$ | SD $F_{IS}$ | Mean Tajima's D | SD Tajima's D |
|---|---|---|---|---|---|---|---|---|---|---|---|
| **Mad** | 0fold | 7958721 | 27976 | 10 | 6.0E-04 | 0.3390 | 9.91E-04 | 0.0839 | 0.2414 | -0.2883 | 1.0804 |
| | 4fold | 1827891 | 24067 | | 1.8E-03 | | 3.71E-03 | 0.0843 | 0.2450 | -0.0178 | 1.1800 |
| **Ger** | 0fold | 7962706 | 25573 | 25 | 4.9E-04 | 0.2607 | 7.17E-04 | 0.8478 | 0.2379 | 1.0536 | 1.1416 |
| | 4fold | 1828777 | 25992 | | 1.9E-03 | | 3.17E-03 | 0.8477 | 0.2377 | 1.1299 | 1.1978 |
| **Rom1** | 0fold | 7961836 | 28653 | 15 | 4.7E-04 | 0.2932 | 9.08E-04 | 0.9029 | 0.2086 | 0.4064 | 1.1119 |
| | 4fold | 1828577 | 30119 | | 1.6E-03 | | 4.16E-03 | 0.9021 | 0.2083 | 0.4607 | 1.1694 |
| **Rom2** | 0fold | 7959849 | 29165 | 15 | 5.7E-04 | 0.2551 | 9.25E-04 | 0.7915 | 0.3618 | 0.3637 | 1.1622 |
| | 4fold | 1828154 | 29560 | | 2.2E-03 | | 4.08E-03 | 0.7985 | 0.3542 | 0.4860 | 1.2621 |
| **Ukr** | 0fold | 7958707 | 24186 | 15 | 3.8E-04 | 0.3152 | 7.67E-04 | 0.8410 | 0.2913 | 1.2689 | 0.8769 |
| | 4fold | 1827806 | 25161 | | 1.2E-03 | | 3.47E-03 | 0.8441 | 0.2811 | 1.3251 | 0.9136 |
| **Gre1** | 0fold | 7963149 | 85320 | 18 | 1.5E-03 | 0.2873 | 2.58E-03 | 0.0331 | 0.0406 | -0.3095 | 0.8987 |
| | 4fold | 1828870 | 78876 | | 5.1E-03 | | 1.04E-02 | 0.0292 | 0.0383 | 0.0297 | 1.0022 |
| **Gre2** | 0fold | 7962994 | 83729 | 20 | 1.8E-03 | 0.3261 | 2.47E-03 | 0.0255 | 0.0519 | -0.2719 | 0.9185 |
| | 4fold | 1828835 | 78819 | | 5.6E-03 | | 1.01E-02 | 0.0181 | 0.0538 | 0.0779 | 1.0045 |
| **Swe** | 0fold | 7963098 | 19239 | 20 | 9.81E-06 | 0.8918 | 5.68E-04 | 0.8277 | 0.2599 | -0.4020 | 0.7551 |
| | 4fold | 1828852 | 19440 | | 1.10E-05 | | 2.50E-03 | 0.8651 | 0.2788 | -0.4638 | 0.6198 |
| **Fra1** | 0fold | 7961749 | 22056 | 20 | 2.9E-04 | 0.3135 | 6.51E-04 | 0.9617 | 0.0853 | -0.1966 | 0.5232 |
| | 4fold | 1828585 | 22344 | | 9.3E-04 | | 2.87E-03 | 0.9655 | 0.0789 | -0.2300 | 0.5620 |
| **Ita1** | 0fold | 7963199 | 47288 | 23 | 1.2E-03 | 0.2778 | 1.35E-03 | 0.0439 | 0.2004 | 0.2808 | 1.1008 |
| | 4fold | 1828878 | 47154 | | 4.3E-03 | | 5.87E-03 | 0.0418 | 0.1851 | 0.4134 | 1.1581 |
| **Ita2** | 0fold | 7963197 | 53047 | 18 | 1.4E-03 | 0.2827 | 1.61E-03 | 0.0710 | 0.1598 | 0.2329 | 1.0229 |
| | 4fold | 1828877 | 52012 | | 5.1E-03 | | 6.86E-03 | 0.0724 | 0.1547 | 0.4410 | 1.0917 |
| **Nor** | 0fold | 7962937 | 18977 | 19 | 3.5E-06 | 0.2616 | 5.54E-04 | 0.8944 | 0.1735 | 0.7220 | 1.1571 |
| | 4fold | 1828810 | 19403 | | 1.3E-05 | | 2.47E-03 | 0.9580 | 0.1182 | 1.0465 | 1.2634 |
| **Fra2** | 0fold | 7960343 | 22649 | 10 | 7.4E-04 | 0.2928 | 8.02E-04 | 0.8344 | 0.3478 | 1.6475 | 0.6322 |
| | 4fold | 1828221 | 22725 | | 2.5E-03 | | 3.50E-03 | 0.8261 | 0.3674 | 1.7136 | 0.6703 |

**Table 2.** Estimates of genetic diversity and divergence for the different gene sets for 0 and 4-fold degenerate sites using genetic data from population Gre2.

| Tissue | No. genes | Degeneracy | No. Sites | No. SNPs | $\pi$ | $\pi_N/\pi_S$ | Watterson's θ | Tajima's D | *d* | $d_N/d_S$ |
|---|---|---|---|---|---|---|---|---|---|---|
| **Male gametophyte** | 12398 | 0fold | 2659597 | 18755 | 9.37E-04 | 1.73E-01 | 1.66E-03 | -3.37E-01 | 4.24E-03 | 0.1490 |
| | | 4fold | 544667 | 19122 | 5.40E-03 | | 8.25E-03 | 5.70E-02 | 2.85E-02 | |
| **Female gametophyte** | 10893 | 0fold | 902454 | 6132 | 9.28E-04 | 1.58E-01 | 1.60E-03 | -2.60E-01 | 4.01E-03 | 0.1320 |
| | | 4fold | 185058 | 6789 | 5.89E-03 | | 8.62E-03 | 1.20E-01 | 3.04E-02 | |
| **Sporophyte (randomly selected)** | 10000 | 0fold | 4116131 | 38473 | 1.67E-03 | 1.81E-01 | 2.20E-03 | -2.53E-01 | 4.86E-03 | 0.1640 |
| | | 4fold | 951241 | 42264 | 9.17E-03 | | 1.04E-02 | 9.38E-02 | 2.96E-02 | |
| **Vegetative pollen** | 4148 | 0fold | 1681068 | 10636 | 8.61E-04 | 1.59E-01 | 1.49E-03 | -2.85E-01 | 3.95E-03 | 0.1375 |
| | | 4fold | 343068 | 12063 | 5.41E-03 | | 8.27E-03 | 5.21E-02 | 2.87E-02 | |
| **Pollen tube** | 5032 | 0fold | 2128017 | 14459 | 9.44E-04 | 1.76E-01 | 1.60E-03 | -2.90E-01 | 4.24E-03 | 0.1500 |
| | | 4fold | 435798 | 14979 | 5.37E-03 | | 8.08E-03 | 6.06E-02 | 2.83E-02 | |
| **Sperm** | 3218 | 0fold | 1412757 | 10425 | 1.02E-03 | 1.87E-01 | 1.73E-03 | -2.89E-01 | 4.60E-03 | 0.1616 |
| | | 4fold | 288647 | 9947 | 5.44E-03 | | 8.10E-03 | 3.98E-02 | 2.85E-02 | |
| **Egg** | 1607 | 0fold | 585660 | 3852 | 9.28E-04 | 1.56E-01 | 1.55E-03 | -2.49E-01 | 4.04E-03 | 0.1364 |
| | | 4fold | 119761 | 4346 | 5.93E-03 | | 8.53E-03 | 1.01E-01 | 2.96E-02 | |
| **Synergids** | 511 | 0fold | 135298 | 817 | 7.75E-04 | 5.67E-03 | 1.42E-03 | -2.53E-01 | 3.55E-03 | 0.1174 |
| | | 4fold | 27399 | 999 | 5.67E-03 | | 8.57E-03 | 1.35E-01 | 3.02E-02 | |
| **Nucellus** | 6875 | 0fold | 2693937 | 18742 | 9.72E-04 | 1.70E-01 | 1.64E-03 | -2.89E-01 | 4.29E-03 | 0.1458 |
| | | 4fold | 554349 | 19738 | 5.72E-03 | | 8.37E-03 | 8.75E-02 | 2.94E-02 | |
| **Unpollinated pistil** | 9555 | 0fold | 4007686 | 35755 | 1.03E-03 | 1.82E-01 | 2.10E-03 | -4.20E-01 | 4.54E-03 | 0.1520 |
| | | 4fold | 928126 | 41465 | 5.64E-03 | | 1.05E-02 | 3.02E-03 | 2.99E-02 | |

**Table 3.** Origin of 228 samples included in this study from 13 different populations.

| Population | No. Samples | Latitude | Longitude |
|---|---|---|---|
| **Mad** | 10 | 32.74 | -16.93 |
| **Ger** | 25 | 49.74 | 11.43 |
| **Rom1** | 15 | 45.47 | 25.23 |
| **Rom2** | 15 | 45.60 | 24.62 |
| **Ukr** | 15 | 48.19 | 24.57 |
| **Gre1** | 18 | 37.96 | 22.41 |
| **Gre2** | 20 | 39.86 | 20.78 |
| **Swe** | 20 | 68.35 | 18.74 |
| **Fra1** | 20 | 44.07 | 7.52 |
| **Ita1** | 23 | 42.50 | 13.58 |
| **Ita2** | 18 | 42.25 | 13.32 |
| **Nor** | 19 | 62.86 | 11.78 |
| **Fra2** | 10 | 45.16 | 5.63 |

**Table 4.** Number of SNPs and invariant sites after applying each filtering step.

| Criteria for filtering | Invariant sites (bp) | SNPs (bp) |
|---|---|---|
| **Unfiltered** | 219,544,869 | 34,243,876 |
| **Missing data (10%)** | 126,912,788 | 19,420,215 |
| **Repetitive regions** | 78,822,573 | 7,721,792 |
| **Coverage** | 34,790,202 | 2,785,366 |
| **Allele balance** | 34,790,202 | 2,776,972 |

**Table 5.** Publicly available ArrayExpress datasets available on EMBL EBI used to characterize genes expressed on each tissue and/or cellular type.

| Tissue/ cellular type | Ploidy | Accession number raw dataset | Reference |
|---|---|---|---|
| **Mature vegetative pollen** | Haploid | E-ATMX-35 | Borges et al. 2008 |
| **Sperm** | Haploid | E-ATMX-35 | Borges et al. 2008 |
| **Pollen tube (semi-*in vivo*)** | Haploid | E-GEOD-17343 | Qin et al. 2009 |
| **Synergid cells** | Haploid | E-MEXP-2227 | Wuest et al. 2010 |
| **Egg** | Haploid | E-MEXP-2227 | Wuest et al. 2010 |
| **Central cell** | Haploid | E-MEXP-2227 | Wuest et al. 2010 |
| **Nucellus** | Diploid | E-MEXP-3138 | Schmid et al. 2012 |
| **Unpollinated pistil** | Diploid | E-GEOD-27281 | Boavida et al. 2011 |

**Table 6.** Publicly available ArrayExpress datasets used to perform expression level correction on data sets.

| Tissue/ cellular type | Accession number raw dataset | Reference |
|---|---|---|
| **Petiole** | E-AFMX-9: ATGE_19 | Schmid et al. 2005 |
| **Root** | E-AFMX-9: ATGE_9 | Schmid et al. 2005 |
| **Flowers stage 12, sepals** | E-AFMX-9: ATGE_34 | Schmid et al. 2005 |
| **Shoot apex, vegetative + young leaves** | E-AFMX-9: ATGE_40 | Schmid et al. 2005 |
| **Pedicels** | E-AFMX-9: ATGE_41 | Schmid et al. 2005 |
| **Leaf** | E-AFMX-9: ATGE_92 | Schmid et al. 2005 |
| **Mature vegetative pollen** | E-ATMX-35 | Borges et al. 2008 |
| **Sperm** | E-ATMX-35 | Borges et al. 2008 |
| **Pollen tube (semi-*in vivo*)** | E-GEOD-17343 | Qin et al. 2009 |
| **Synergid cells** | E-MEXP-2227 | Wuest et al. 2010 |
| **Egg** | E-MEXP-2227 | Wuest et al. 2010 |
| **Central cell** | E-MEXP-2227 | Wuest et al. 2010 |
| **Nucellus** | E-MEXP-3138 | Schmid et al. 2012 |

### Figures

**Figure 1.** Map depicting 4-fold $\pi$ values for each population.

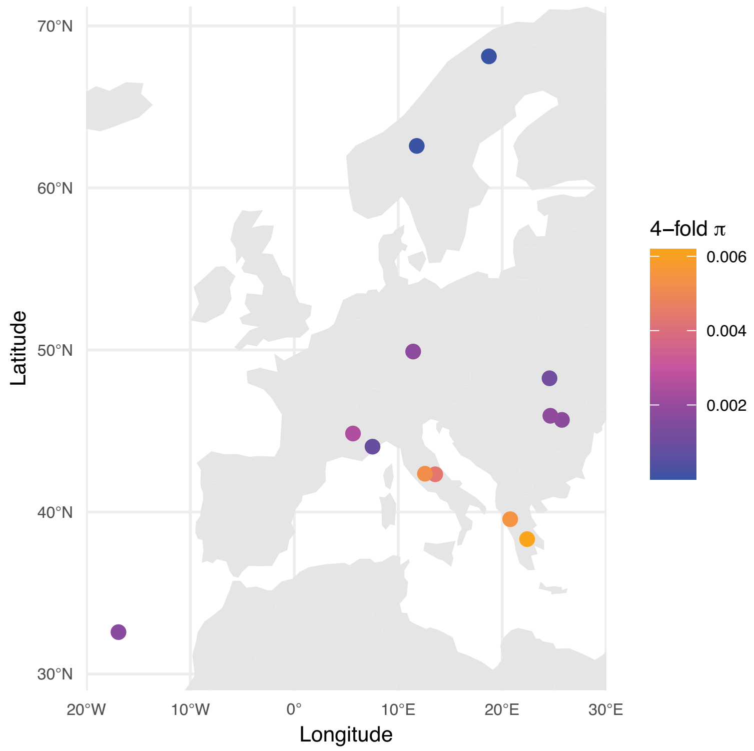

**Figure 2.** $F_{IS}$ values for 0-fold degenerate and 4-fold degenerate SNPs.

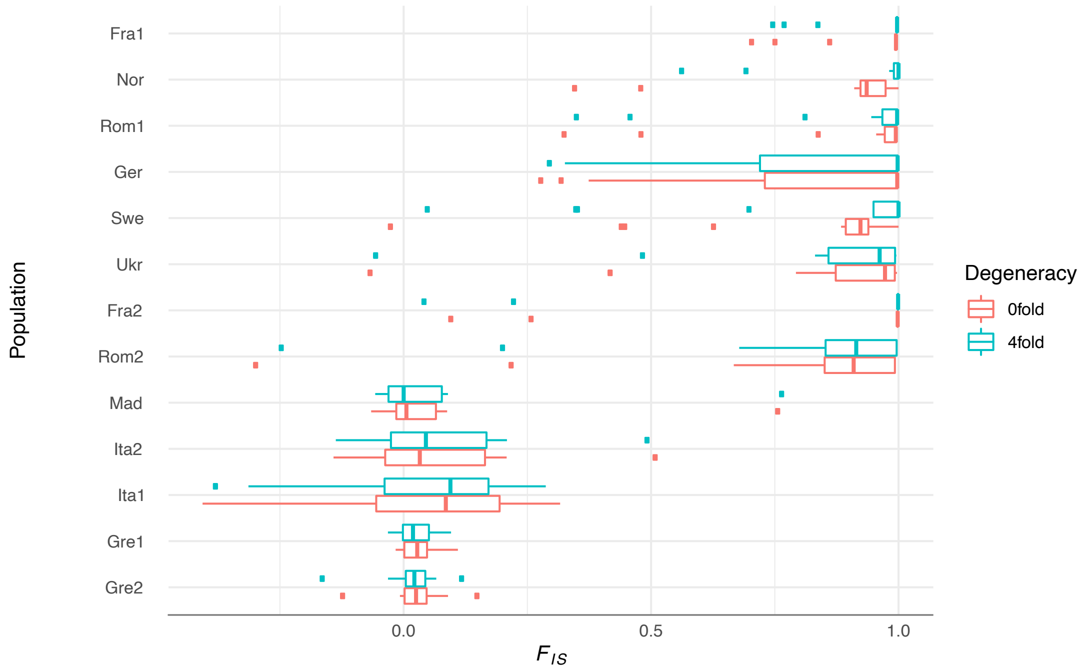

**Figure 3.** Estimates of the proportion of the genome in runs of homozygosity above 500 Kb ($F_{ROH}$).

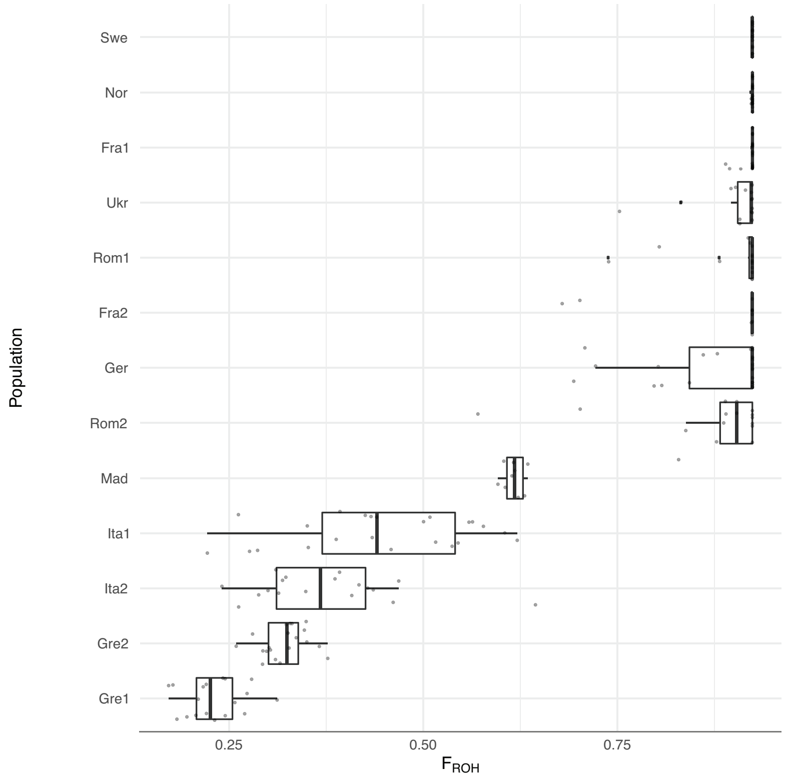

**Figure 4.** (**A**) Proportion of adaptive substitutions (*α*) and (**B**) the proportion of adaptive substitutions relative to neutral divergence ($\omega_\alpha$) between the male and female gametophyte male gametophyte (*n*=12,398), female gametophyte (*n*=10,893) and sporophytic tissues (*n*=10,000, random sample) from population Gre2. We identified significant ($P<0.05$) differences between groups (indicated by different letters) based on a Kruskal-Wallis test followed by a post-hoc Dunn test with Bonferroni correction. Error bars represent the the bootstrap standard error (200 replicates).

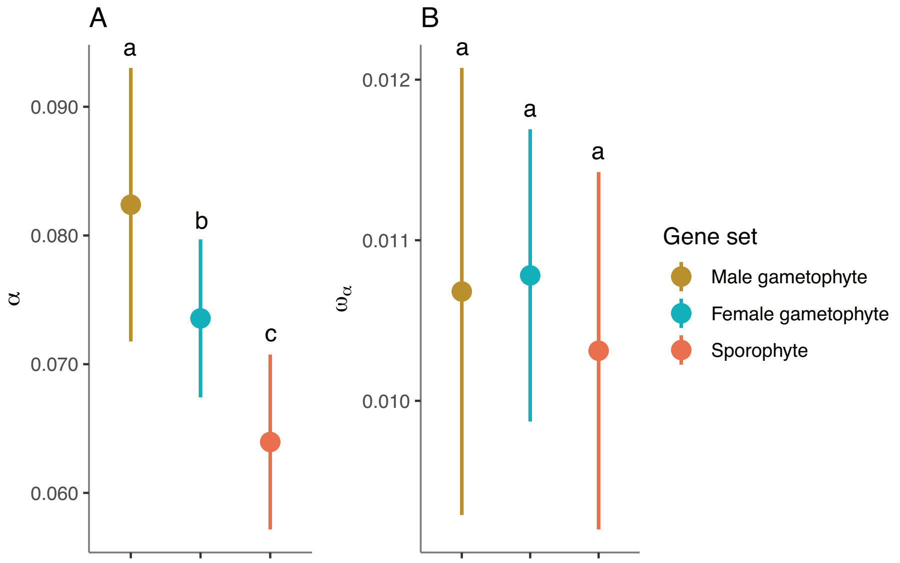

**Figure 5.** Proportion of adaptive substitutions relative to neutral divergence ($\omega_\alpha$) (**A**) Genes expressed in vegetative pollen (*n*=4,148), sperm (*n*=3,218), pollen tubes (*n*=5,032), synergids (*n*=511), egg (*n*=1,607), nucellus (*n*=6,875) and unpollinated pistils (*n*= 9,555). (**B**) Genes exclusively expressed in vegetative pollen (*n*=144), pollen tubes (*n*=309) and sperm cells (*n*=260). (**C**) Genes matching a common distribution of expression levels: vegetative pollen (*n*=1,552), pollen tubes (*n*=2,387) and sperm (*n*=3,218). Error bars represent the bootstrap standard error (200 replicates). Different letters denote statistically significant differences between gene set categories ($P$<0.05). We identified significant (P<0.05) differences between groups (indicated by different letters) based on a Kruskal-Wallis test followed by a post-hoc Dunn test with Bonferroni correction.

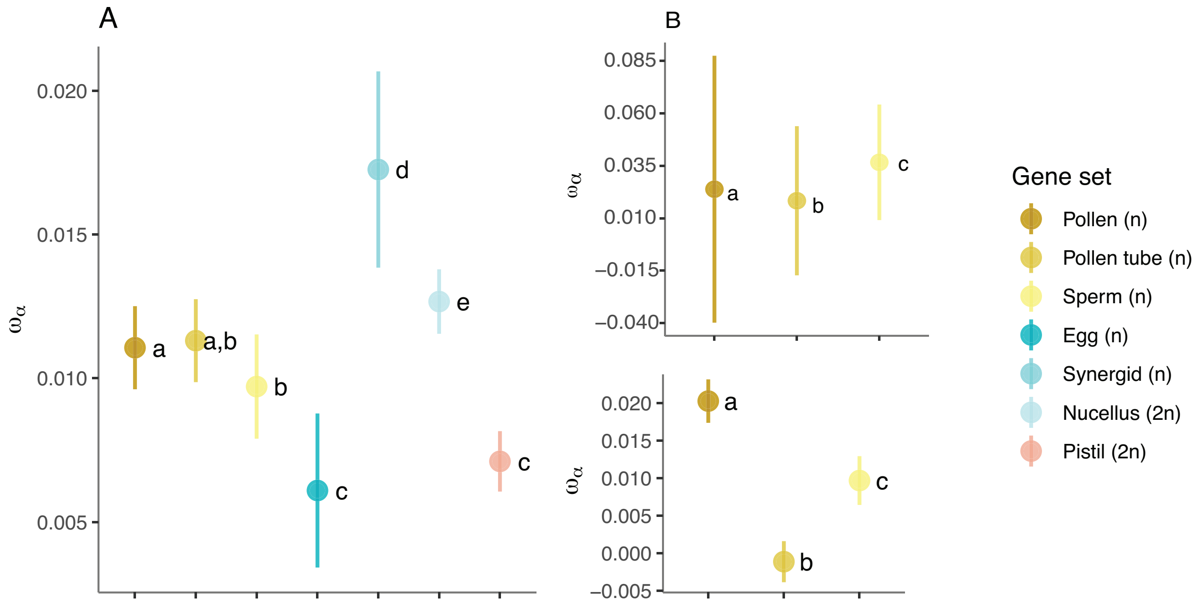

**Figure 6.** Comparison of (**A**) DFE estimates, (**B**) the proportion of adaptive substitutions ($\alpha$) and (**C**) the proportion of adaptive substitutions relative to neutral divergence ($\omega_\alpha$) between sets of genes expressed in pollen and unpollinated pistils using expression data from *Arabidopsis thaliana* and *Brassica napus* for population Gre2. Data set sizes: pollen-expressed genes in *A. thaliana*=4,148; pollen-expressed genes in *B. napus*=2,489; pistil-expressed genes in *A. thaliana*=9,555; pistil-expressed genes in *B. napus*=9,162. Error bars represent the bootstrap standard error (200 replicates). Different letters denote statistically significant differences between gene set categories (*P*<0.05). We identified significant (P<0.05) differences between groups (indicated by different letters) based on a Kruskal-Wallis test followed by a post-hoc Dunn test with Bonferroni correction.

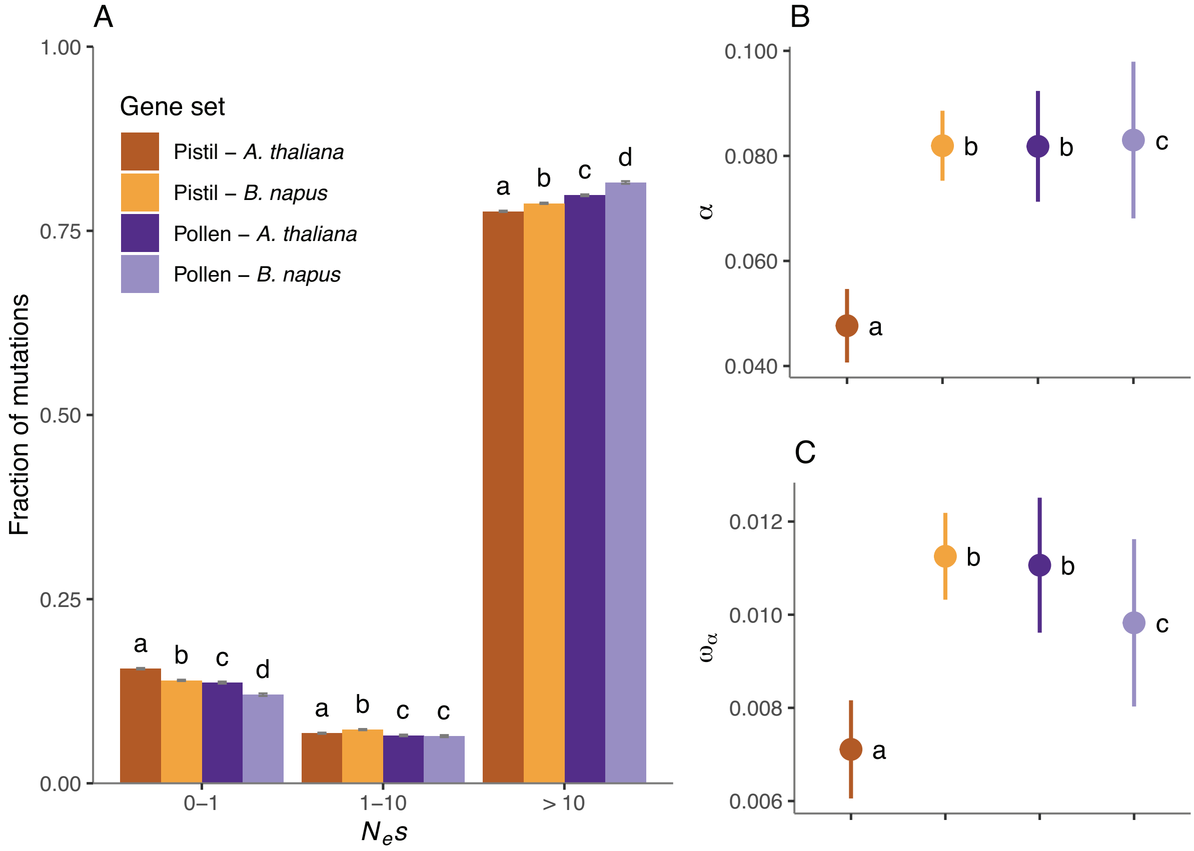

**Figure 7.** Comparison of (**A, B**) DFE estimates, (**C, D**) the proportion of adaptive substitutions ($\alpha$) and (**E, F**) the proportion of adaptive substitutions relative to neutral divergence ($\omega_\alpha$) Gre2, (**A, C, E**) for sets of genes putatively exclusive to vegetative pollen (*n*=144), pollen tubes (*n*=309), sperm cells (*n*=260) and the sporophyte (*n*=5,721). (**B, D, F**) We repeated these analyses using sets of genes identified as exclusively expressed by intersecting each set with genes expressed in seedlings (vegetative pollen= 93, pollen tubes=306, sperm=219 and seedlings=5,298 genes). Error bars represent the bootstrap standard error (200 replicates). Different letters denote statistically significant differences between gene set categories (*P*<0.05). We identified significant (P<0.05) differences between groups (indicated by different letters) based on a Kruskal-Wallis test followed by a post-hoc Dunn test with Bonferroni correction.

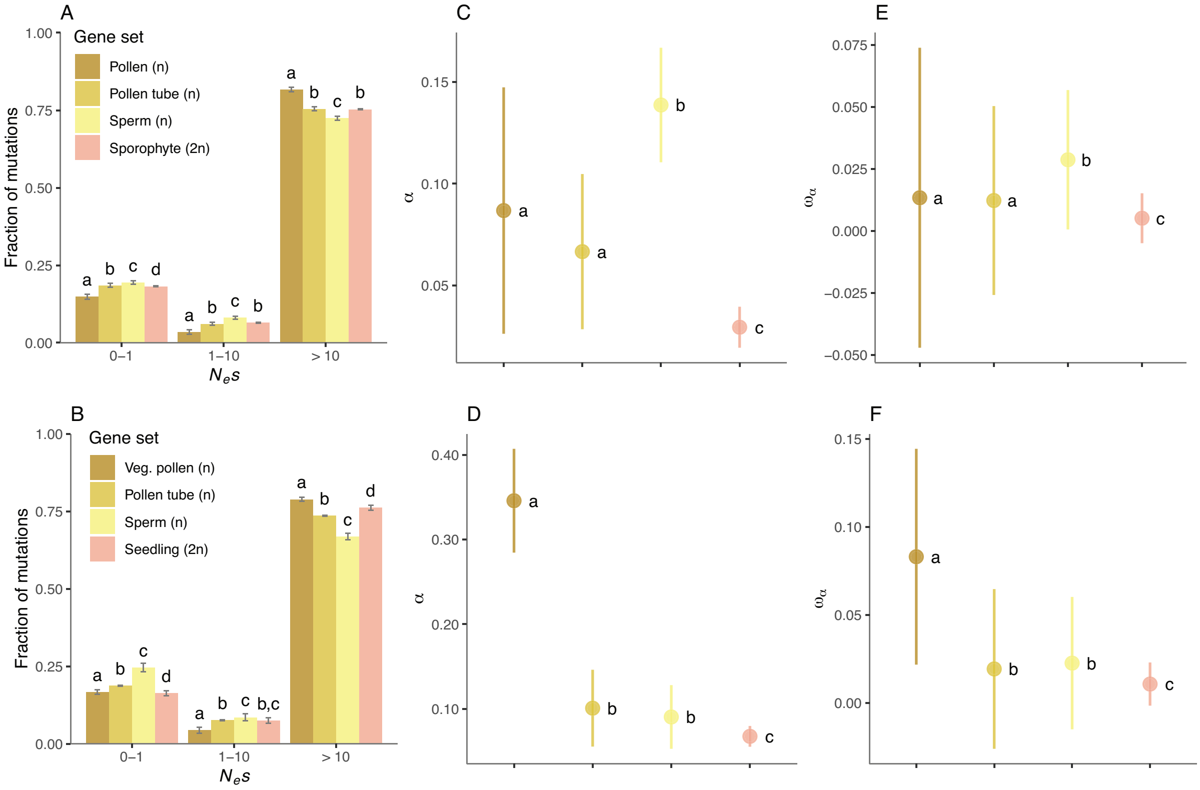

**Figure 8.** Frequency polygons depicting the proportional distribution of expression level values (based on RMA estimates, see Materials and Methods: *Controlling for Gene Expression Breadth and Transcript Abundance*) for each gene set before and after fitting to the distribution of sperm-expressed genes (*n*=3,218; median=824). (**A**) The set of pollen-expressed genes (*n*=4,148; median=1,092) was reduced to 37% of the original list (*n*=1,552; median=837), and (**B**) the set of pollen tubes-expressed genes (*n*=5,032; median=862) was decreased to 47% of the initial data set (*n*=2,387; median correction=844).

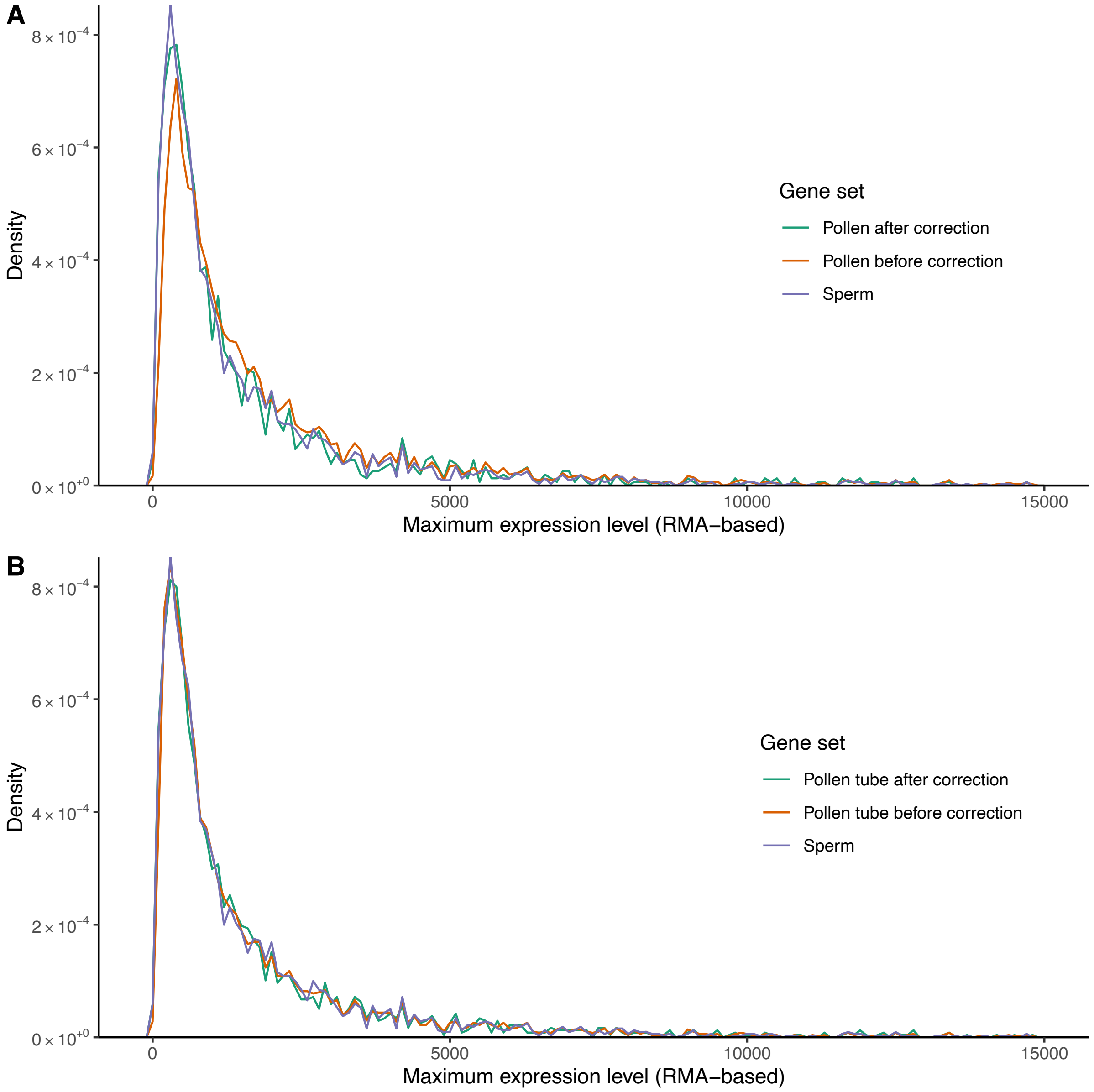

**Figure 9.** Assignment of individuals to each of the inferred ancestral population clusters largely coincides with samples's geographic origin. (**A**) Geographic origin of 13 populations of *A. alpina* included in this study. (**B**) Bayesian clustering of 228 individuals showing ancestral proportions for *K*=8 clusters based on fastSTRUCTURE results.

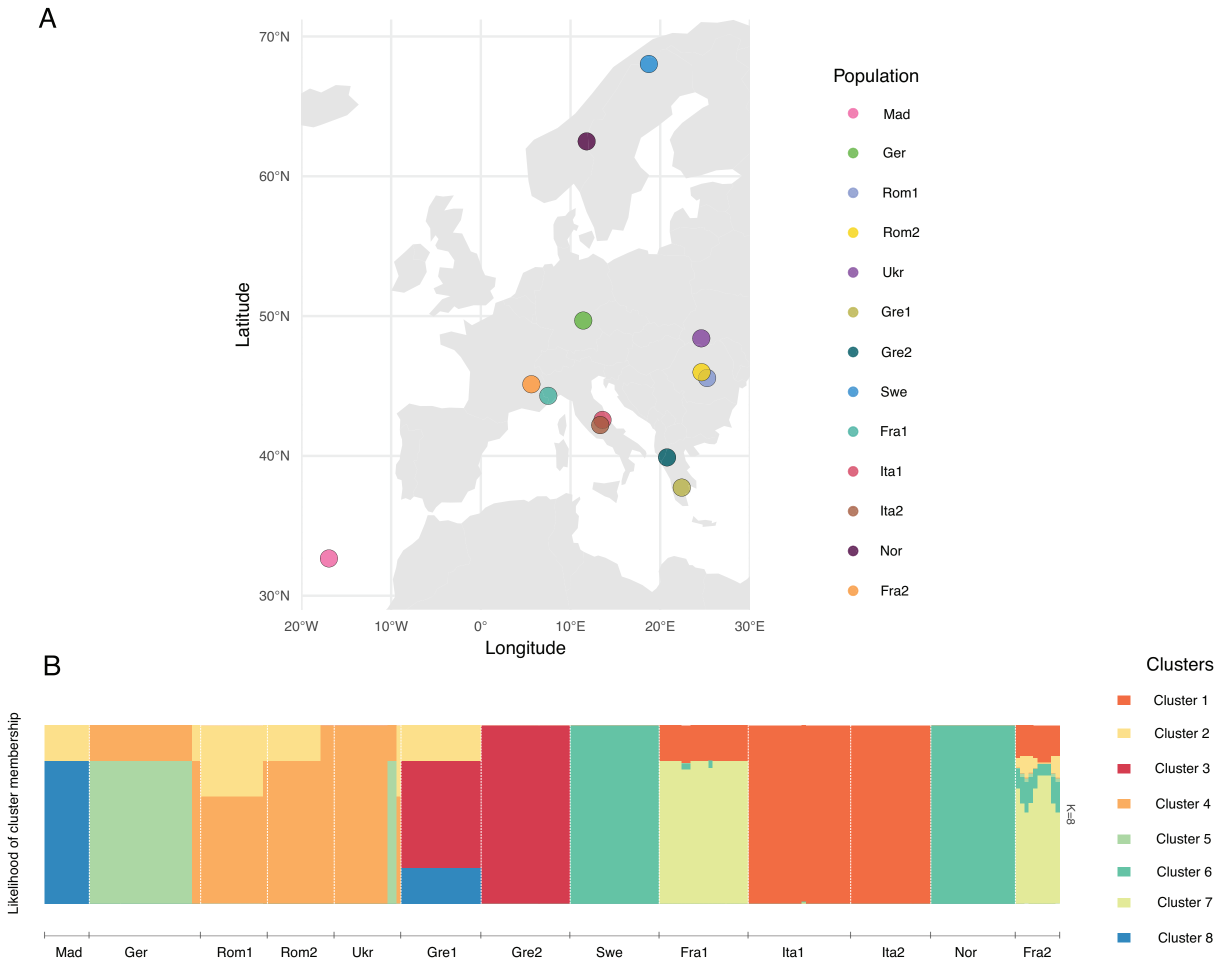

**Figure 10.** Scatterplots depicting the correlation between (**A-D**) $F_{ROH}$, (**E-H**) 4-fold $\pi$, and (**I-L**) mean 4-fold $F_{IS}$ respect to the fraction of nearly neutral sites ($0 < N_e s < 1$) for vegetative pollen, pollen tubes, sperm and a set of 10,000 randomly selected genes used as a reference of genome-wide estimates.

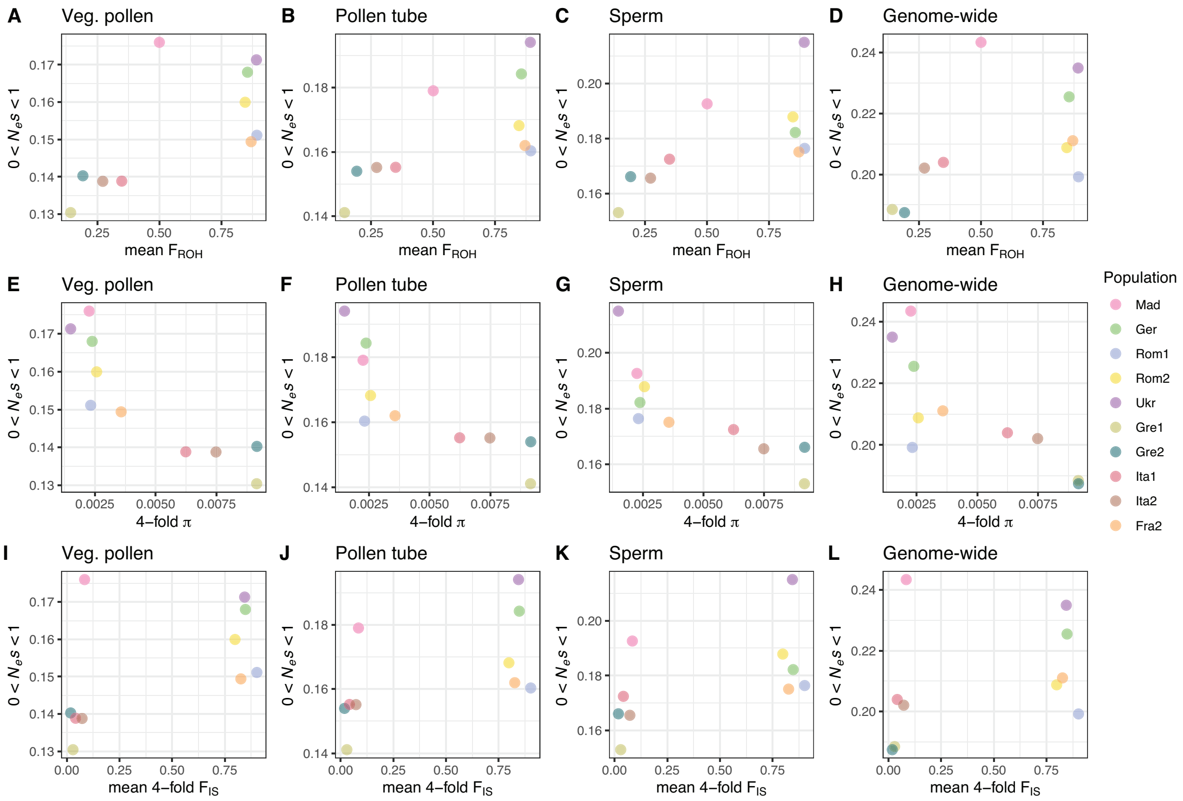

**Figure 11.** Inferred DFE for 10,000 randomly selected genome-wide genes for all populations. Populations Swe, Nor, and Fra1 show genome-wide relaxed purifying selection (i.e., the majority of mutations experience weak to mild purifying selection, $N_e s < 10$).

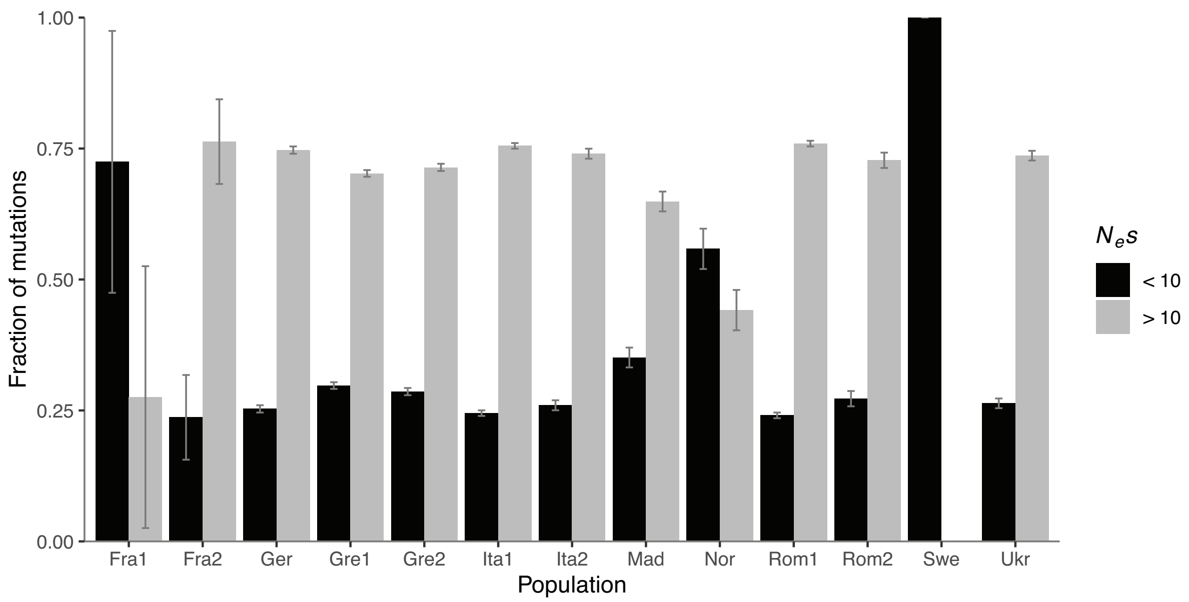

**Figure 12.** Scatterplots depicting the correlation between PIC-corrected estimates of mean 4-fold $\pi$ and the fraction of nearly neutral sites ($0 < N_e s < 1$). Spearman's rank correlation rho ($\rho$) values and the statistical significance of the association between these variables are shown for (**A**) vegetative pollen, (**B**) pollen tubes, (**C**) sperm and (**D**) a set of 10,000 randomly selected genes genome-wide (*: $P$<0.05, **: $P$<0.01, ***: $P$<0.001).

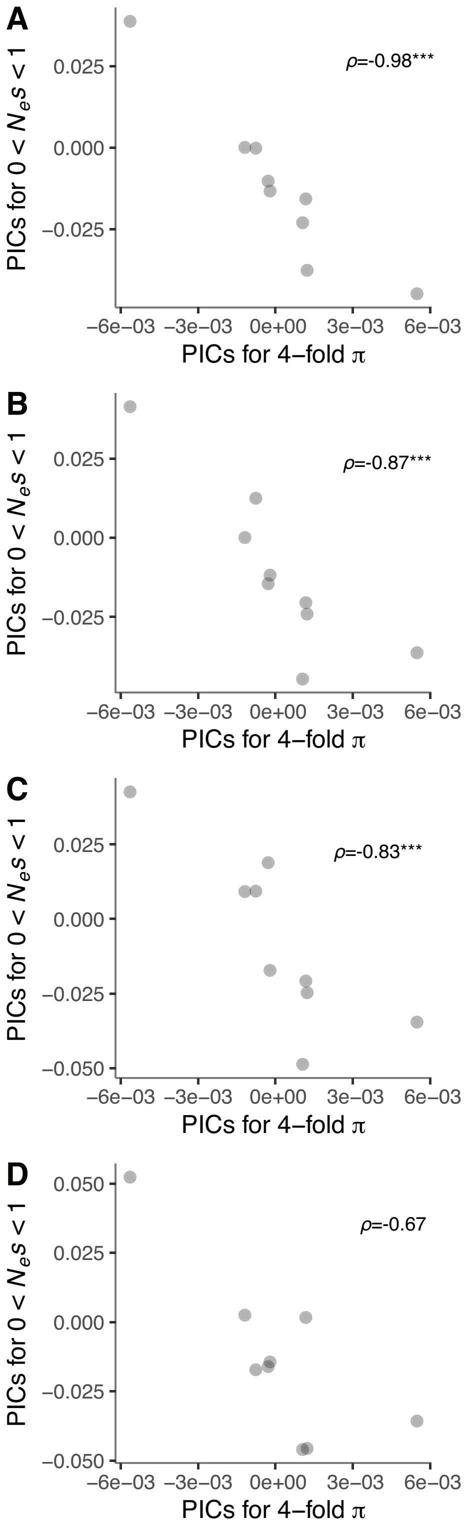

**Figure 13.** Estimates of the fraction of new nearly neutral nonsynonymous mutations, ($0 < N_e s < 1$), across populations for sets of genes expressed in vegetative pollen (*n*=4,148), sperm (*n*=3,218) and genome-wide estimates (*n*=10,000, randomly selected genes). Error bars represent the bootstrap standard error (200 replicates). Different letters denote statistically significant differences between gene set categories ($P$<0.05). We identified significant (P<0.05) differences between groups (indicated by different letters) based on a Kruskal-Wallis test followed by a post-hoc Dunn test with Bonferroni correction.

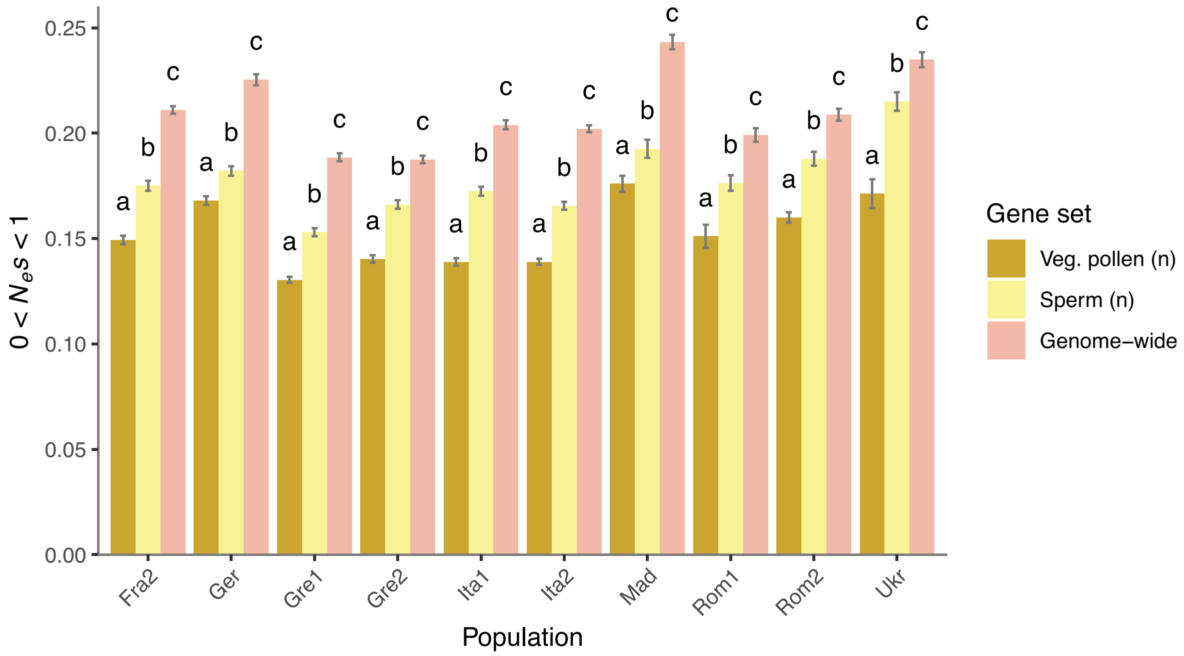

**Figure 14.** Scatterplot depicting the correlation between estimates of 4-fold *π* and the fraction of nearly neutral sites (0 < $N_e s$ < 1) for genes expressed in the synergids across *A. alpina* populations (n=10) (**A**) before and (**B**) after correction for phylogenetic non-independence (NS in both cases: *P*>0.05).

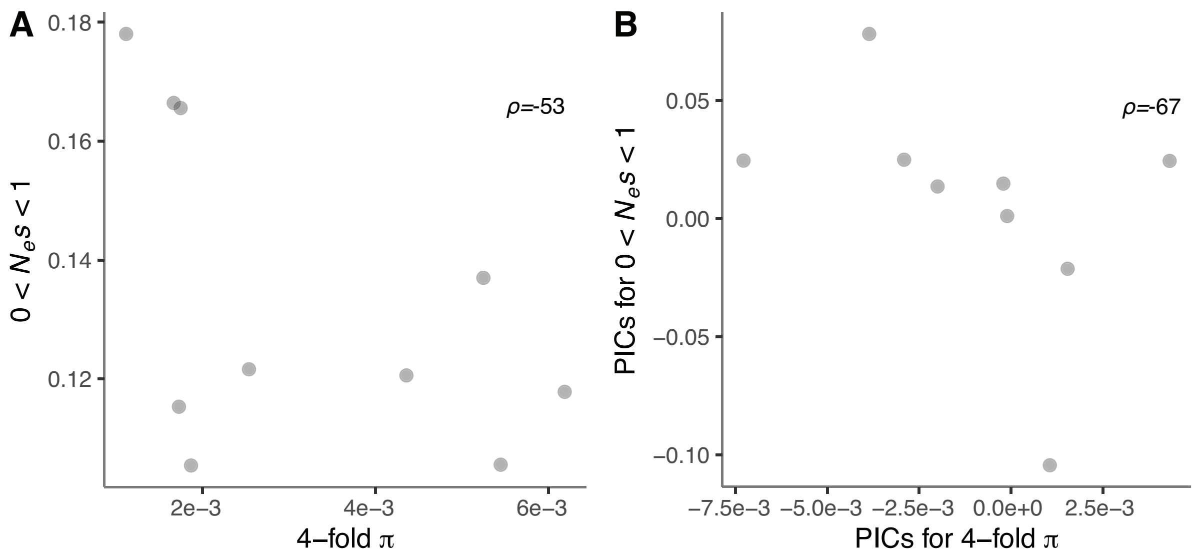

**Figure 15.** (**A**) ML phylogeny of ten populations of *A. alpina* (*n*=169 samples) based on 15,033 markers. (**B**) Reduced ML tree obtained after randomly subsampling each population to a single tip. Numbers at nodes indicate bootstrap support (n=500 replicates). Scale bars indicate the number of nucleotide substitutions per site.

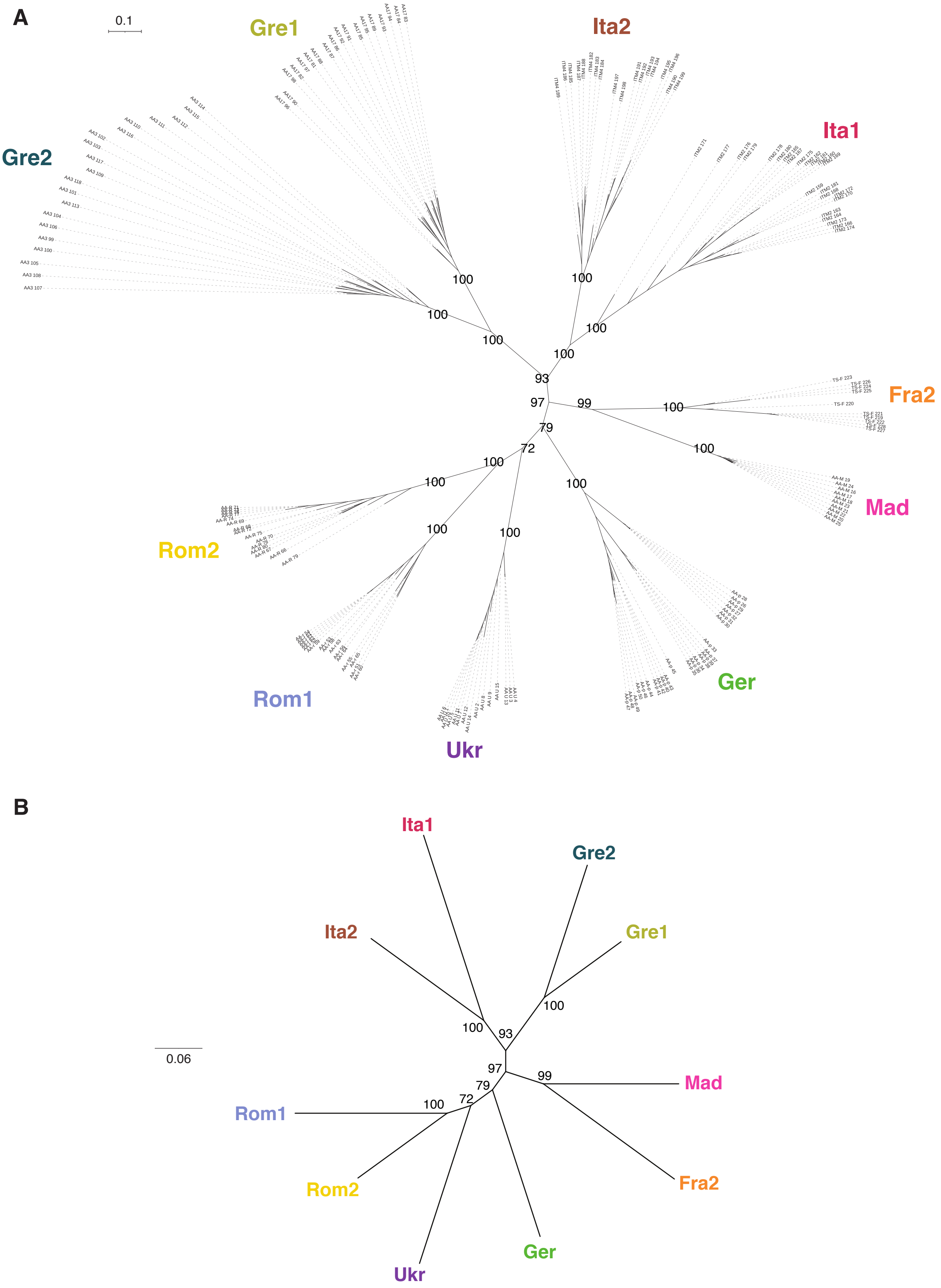

### References

Arunkumar R, Josephs EB, Williamson RJ, Wright SI. 2013. Pollen-Specific, but not Sperm-Specific, Genes Show Stronger Purifying Selection and Higher Rates of Positive Selection than Sporophytic Genes in Capsella grandiflora. Mol Biol Evol. 30:2475-2486.

Boavida LC, Borges F, Becker JD, Feijó JA. 2011. Whole Genome Analysis of Gene Expression Reveals Coordinated Activation of Signaling and Metabolic Pathways During Pollen-Pistil Interactions in *Arabidopsis*. Plant Physiol. 155:2066-80.

Bolger AM, Lohse M, Usadel B. 2014. Trimmomatic: a Flexible Trimmer for Illumina Sequence Data. Bioinformatics. 30:2114-2120.

Borges F, Gomes G, Gardner R, Moreno N, McCormick S, Feijó JA, Becker JD. 2008. Comparative Transcriptomics of *Arabidopsis* Sperm Cells. Plant Physiol. 148:1168-1181.

Broad Institute. 2019. Picard Tools. Broad Institute, GitHub Repository. http://broadinstitute.github.io/picard/

Chalhoub B, Denoeud F, Liu S, Parkin IA, Tang H, Wang X, Chiquet J, Belcram H, Tong C, Samans B, et al. 2014. Early Allopolyploid Evolution in the Post-Neolithic Brassica napus Oilseed Genome. Science. 345:950-3.

Jiao W-B, Accinelli GG, Hartwig B, Kiefer C, Baker D, Severing E, Willing E-M, Piednoel M, Woetzel S, Madrid-Herrero E, et al. 2017. Improving and Correcting the Contiguity of Long-Read Genome Assemblies of Three Plant Species Using Optical Mapping and Chromosome Conformation Capture Data. Genome Res. 27:778-786.

Li H, Durbin R. 2009. Fast and Accurate Short Read Alignment with Burrows-Wheeler Transform. Bioinformatics. 25:1754-1760.

Lohani N, Singh MB, Bhalla PL. 2021. RNA-Seq Highlights Molecular Events Associated with Impaired Pollen-Pistil Interactions Following Short-Term Heat Stress in Brassica napus. Front Plant Sci. 11:622748.

Martin M. 2011. Cutadapt Removes Adapter Sequences from High-Throughput Sequencing Reads. EMBnet.journal. 17:10-12.

Mammana A, Helmuth, J. 2020. bamsignals: Extract Read Count Signals from Bam Files. https://github.com/lamortenera/bamsignals.

McKenna A, Hanna M, Banks E, Sivachenko A, Cibulskis K, Kernytsky A, Garimella K, Altshuler D, Gabriel S, Daly M, et al. 2010. The Genome Analysis Toolkit: a MapReduce framework for analyzing next-generation DNA sequencing data. Genome Res. 20:1297-1303.

R Core Team. 2019. R: A language and environment for statistical computing. R Foundation for Statistical Computing. Vienna, Austria. URL https://www.R-project.org/.

Poplin R, Ruano-Rubio V, DePristo MA, Fennell TJ, Carneiro MO, Van der Auwera GA, Kling DE, Gauthier LD, Levy-Moonshine A, Roazen D, et al. 2017. Scaling Accurate Genetic Variant Discovery to Tens of Thousands of Samples. bioRxiv:201178.

Qin Y, Leydon AR, Manziello A, Pandey R, Mount D, Denic S, Vasic B, Johnson MA, Palanivelu R. 2009. Penetration of the Stigma and Style Elicits a Novel Transcriptome in Pollen Tubes, Pointing to Genes Critical for Growth in a Pistil. PLoS Genet. 5:e1000621.

Schmid M, Davison TS, Henz SR, Pape UJ, Demar M, Vingron M, Schölkopf B, Weigel D, Lohmann JU. 2005. A Gene Expression Map of *Arabidopsis thaliana* Development. Nat Genet. 37:501-506.

Schmid MW, Schmidt A, Klostermeier UC, Barann M, Rosenstiel P, Grossniklaus U. 2012. A Powerful Method for Transcriptional Profiling of Specific Cell Types in Eukaryotes: Laser-Assisted Microdissection and RNA Sequencing. PLoS ONE. 7:e29685.

Smit AFA, Hubley R, Green P. 2019. RepeatMasker. http://www.repeatmasker.org.

Wagner GP, Kin K, Lynch VJ. 2013. A Model Based Criterion for Gene Expression Calls Using RNA-seq Data. Theory Biosci.132:159-64.

Willing E-M, Rawat V, Mandáková T, Maumus F, James GV, Nordström KJV, Becker C, Warthmann N, Chica C, Szarzynska B, et al. 2015. Genome Expansion of *Arabis Alpina* Linked with Retrotransposition and Reduced Symmetric DNA Methylation. Nat Plants. 1:14023.

Wuest SE, Vijverberg K, Schmidt A, Weiss M, Gheyselinck J, Lohr M, Wellmer F, Rahnenführer J, von Mering C, Grossniklaus U. 2010. *Arabidopsis* Female Gametophyte Gene Expression Map Reveals Similarities Between Plant and Animal Gametes. Curr Biol. 20:506-512.
